# The super repertoire of type IV effectors in the pangenome of *Ehrlichia* spp. provides insights into host-specificity and pathogenesis

**DOI:** 10.1101/2020.11.07.372862

**Authors:** Christophe Noroy, Damien F. Meyer

## Abstract

The identification of bacterial effectors is essential to understand how obligatory intracellular bacteria such as *Ehrlichia* spp. manipulate the host cell for survival and replication. Infection of mammals – including humans – by the intracellular pathogenic bacteria *Ehrlichia* spp. depends largely on the injection of virulence proteins that hijack host cell processes. Several hypothetical virulence proteins have been identified in *Ehrlichia* spp., but one so far has been experimentally shown to translocate into host cells via the type IV secretion system. However, the current challenge is to identify most of the type IV effectors (T4Es) to fully understand their role in *Ehrlichia* spp. virulence and host adaptation. Here, we predict the T4E repertoires of four sequenced *Ehrlichia* spp. and four other *Anaplasmataceae* as comparative models (pathogenic *Anaplasma* spp. and *Wolbachia* endosymbiont) using previously developed S4TE 2.0 software. This analysis identified 579 predicted T4Es (228 pT4Es for *Ehrlichia* spp. only). The effector repertoires *of Ehrlichia* spp. overlapped, thereby defining a conserved core effectome of 92 effectors shared by all strains. In addition, 69 species-specific T4Es were predicted with non-canonical GC% mostly in gene sparse regions of the genomes and we observed a bias in pT4Es according to host-specificity. We also identified new protein domain combinations, suggesting novel effector functions. This work presenting the predicted effector collection *of Ehrlichia* spp. can serve as a guide for future functional characterisation of effectors and design of alternative control strategies against these bacteria.

## Introduction

Gram-negative intracellular bacteria *Ehrlichia* spp. are pathogens of eukaryotic cells. They have evolved to survive and replicate in a wide range of tick and mammalian hosts, including humans. Following the first publication of the *Ehrlichia chaffeensis* genome sequence in 2006 (Dunning Hotopp et al., 2006), four other species of pathogenic *Ehrlichia* with a versatile host range have been sequenced (https://gold.jgi.doe.gov/). For instance, *E. chaffeensis*, which is the agent of human monocytic ehrlichiosis, can also cause disease in several other vertebrates, including dogs and deer (Paddock and Childs, 2003) whereas *E. canis, E. muris* and *E. ruminantium* have narrower host ranges (Braga et al., 2014; Feng and Walker, 2004; Peter et al., 2002).

For infection, *Ehrlichia* spp. depends on a dedicated protein complex, the type IV secretion system (T4SS), which acts as a molecular syringe to translocate bacterial proteins into the host cells (Moumène and Meyer, 2016). The study of type IV effectors (T4Es) provides valuable insights into the mechanisms by which an intracellular pathogen can manipulate eukaryotic cellular processes to survive and replicate in host cells (Rikihisa, 2017; O’Connor et al., 2012; Martinez et al., 2016). In Gram-negative intracellular bacteria, a large number of effectors harbour eukaryotic-like domains (Ninio and Roy, 2007). These proteins interfere in different steps of the infection by mimicking the functions of eukaryotic proteins (Cazalet et al., 2004; de Felipe et al., 2008). Being able to predict these eukaryotic domains, as well as other protein-protein interaction motif or subcellular signalling sequences, is an important step towards understanding the effectors’ mode of action. The identification of T4Es, as well as the elucidation of their function inside the host cell, will help understanding the bacterial pathogenesis. A possible approach for the prediction of T4Es is the analysis of T4E protein sequences using machine learning (Wang et al., 2017). For this purpose, our laboratory developed the S4TE2.0 algorithm, which searches for a large number of motifs related to the function (eukaryotic-like domains, protein-protein interaction, *etc*.) and the subcellular location of predicted T4Es (Meyer et al., 2013; Noroy et al., 2018).

In different pathogenic bacteria, some effectors appear to be acquired by pathogenic bacteria from eukaryotic cells by horizontal gene transfer (de Felipe et al., 2005; Lurie-Weinberger et al., 2010; Ruh et al., 2017). Previous comparative effectomics of closely related *Ehrlichia chaffeensis* strains at the intra-species level showed that pT4Es repertoires are strongly conserved in these genomes. Despite this strong conservation and the strong selective pressure due to their obligate intracellular way of life, at least one strain-specific pT4E (ECHLIB_RS02720) has been identified in *E. chaffeensis* str. Liberty. This effector appears to be involved in the differential inter-strain virulence observed between Arkansas and Liberty strains in SCID mice (Noroy and Meyer, 2016). Moreover, some intense recombination events between *E. ruminantium* strains have been discovered, which may facilitate bacterial adaptation for survival and adaptation under various environmental conditions in both vector and host species (Cangi et al., 2016). These findings suggest that genomic plasticity plays an important role in the evolution of these bacteria by horizontal gene transfer and recombination events.

The genomes of many animal and plant pathogenic bacteria have been completely sequenced in recent years. Comparative genomics studies demonstrated that repertoires of virulence-associated genes comprise a conserved and variable set of genes among bacterial species but these genes may have different evolutionary histories and play distinct roles in pathogenicity (Hajri et al., 2009; Burstein et al., 2016). In comparative genomics, it is important to know all bacterial strains and pathovar to fully understand the identification of the molecular determinants of virulence and host specificity (Hajri et al., 2009). In this study, to overcome the problem of missing data on new species of *Ehrlichia*, we crossed results obtained for the genera *Ehrlichia* with those of the outgroup *Anaplasma*, which share several hosts.

Even though a large number of T4SS-translocated proteins have been identified in numerous bacteria, only two T4Es have been functionally characterised in the family *Anaplasmataceae*, and their role in invasion and pathogenesis is crucial. AnkA, was identified in *Anaplasma phagocytophilum*, based on sequence homology with repeated ankyrin motifs (Caturegli et al., 2000). Once secreted by T4SS, AnkA is tyrosine-phosphorylated and then directed into the nucleus of the host cell to silence CYBB gene expression (Park et al., 2004; Lin et al., 2007; Garcia-Garcia et al., 2009). The other known *Anaplasmataceae* effector, Ats-1, was identified in *A. phagocytophilum* and has an orthologue in *E. chaffeensis* (Etf-1) (Niu et al., 2010). Ats-1 is injected by T4SS into the cytoplasm of the host cell to recruit host autophagosomes to the bacterial inclusion. Another portion of Ats-1 targets mitochondria, where it has an antiapoptotic activity (Niu et al., 2012; Liu et al., 2012). In contrast, *Legionella pneumophila* relies on a set of approximately 300 T4Es (10% of the genome) for efficient pathogenesis (Gomez-Valero et al., 2014). This molecular arsenal targets many cell signalling and biochemical pathways in the host cell and allows the bacterium to hijack host immunity to ensure its survival and development. We thus hypothesised that many effectors remain to be identified in the *Ehrlichia* genus.

In this study, we predict the pangenome super-repertoire of T4Es and investigate how they are related to genome plasticity and host specificity. We show that T4E gene repertoires of *Ehrlichia* spp. comprise core and variable gene suites, which probably have distinct roles in pathogenicity and different evolutionary histories. By analysing the protein architecture of these effectors, we explored new functions of interest potentially involved in the pathogenesis of this important genus of zoonotic bacteria. Our work thus provides resources for functional and evolutionary studies aiming at understanding the host specificity of *Anaplasmataceae*, functional redundancy between T4Es and the driving forces shaping T4E repertoires.

## Methods

### Retrieval of genome sequences and prediction of type IV effectors

The complete genome sequences of the eight *Anaplasmataceae* studied were obtained from the National Center for Biotechnology Information (NCBI) database (ftp://ftp.ncbi.nih.gov/genomes/Bacteria/). Of the eight bacteria, four are *Ehrlichia* species: *E. chaffeensis* str. Arkansas (NC_007799.1), *E. canis* str. St Jake (NC_007354.1), *E. muris* AS145 (NC_023063.1) and *E. ruminantium* str. Gardel (NC_006831.1). Three are *Anaplasma* species: *A. phagocytophilum* str. HZ (NC_007797.1), *A. marginale str. Florida (NC_012026.1) and A. centrale str. Israel (NC_013532.1*). One is a *Wolbachia* species: *W. endosymbiont* of *D. melanogaster* (NC_002978.6). The repertoires of predicted type IV effectors (pT4Es) were determined using the S4TE 2.0 algorithm with default parameters (Noroy et al., 2018). S4TE 2.0 predicts and ranks candidate T4Es by using a combination of 11 independent modules to explore 14 characteristics of type IV effectors. One module searches for consensus motifs in promoter regions; three modules search for five canonical features of the type IV secretion signal (C-terminal basicity, C-terminal charges, C-terminal hydrophobicity, overall hydrophilicity, and E-blocks); six modules search for several protein domains in known T4Es (eukaryotic–like domains, DUF domains, EPIYA motifs, nuclear localization signals (NLS), mitochondrial localisation signals (MLS), prenylation domains, coiled-coil domains); and one module searches for global homology with known T4Es (Noroy et al., 2018).

### Phylogenetic reconstruction and plasticity related to putative effectors

An initial evolutionary tree was reconstructed based on alignment of the concatenated core genome of *Anaplasmataceae*. The core genomes of the eight studied bacteria were defined using PanOCT software with the following parameters: E-value 10-5, percent identity ≥ 30, and length of match ≥ 65 (Fouts et al., 2012) and resulted in 554 orthologous genes (Fig. 1). To evaluate whether the core predicted type IV effectomes resulted in strong evolutionary events, the tree was reconstructed based on alignment of five concatenated core effectors. Only orthologous genes present in the eight bacteria were used for this phylogenetic reconstruction (Fig. S1). All tree reconstructions were done using MAFFT multiple alignments with default parameters (Katoh et al., 2017) and RAxML under the GAMMA BLOSUM62 model with 100 bootstrap resamplings (Stamatakis, 2014). *W. endosymbiont of D. melanogaster* was used as an out group to root the tree. The Circos algorithm (Krzywinski et al., 2009) was used to represent effector rearrangements between the four *Ehrlichia* species (Fig. 3). By indicating each homology between pT4Es using a spectral color scheme, patterns can be quickly spotted and interpreted. Homologies between effectors were defined using the S4TE-CG algorithm (S4TE 2.0 comparative genomic tool) (Noroy et al., 2018), and PanOCT software (Fouts et al., 2012), and homologous pT4ES were also plotted on a Venn diagram (Fig. S2).

**Figure 1.**
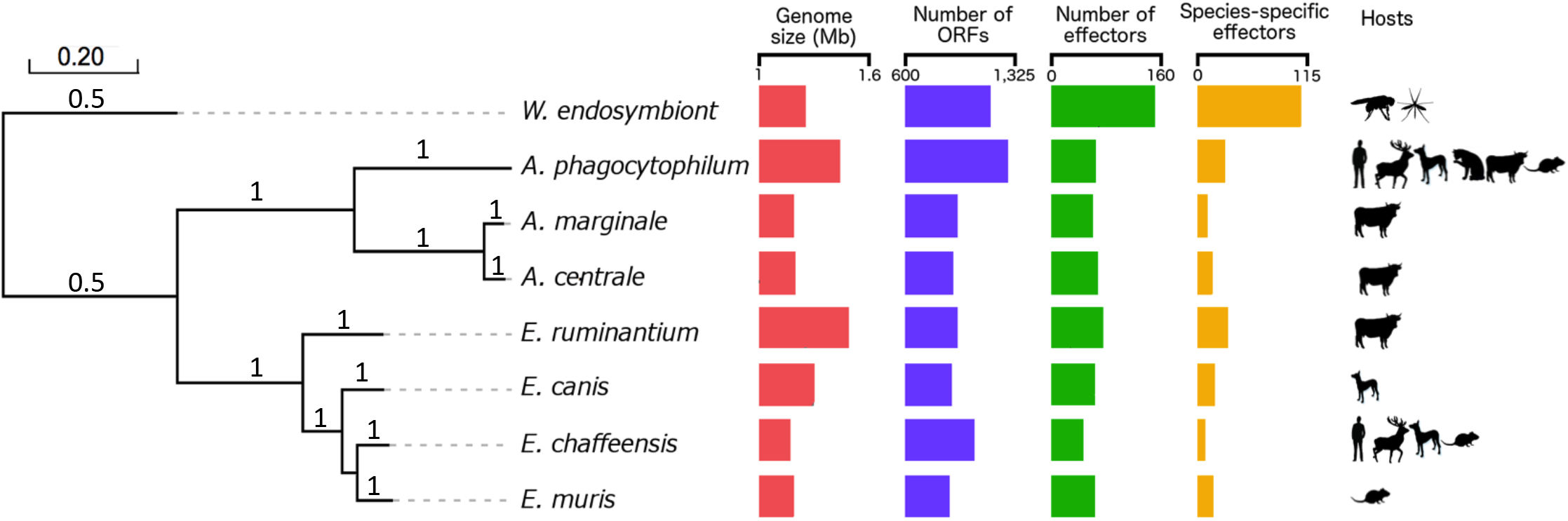
Phylogenetic tree of *Ehrlichia* and *Anaplasma* genus shows three different clades. A maximum likelihood tree of four *Ehrlichia* species *(E. chaffeensis* str. Arkansas, *E. canis* str. Jake, *E. muris* AS145, *E. ruminantium* str. Gardel), three *Anaplasma* species (*A. phagocytophilum* str. HZ, *A. marginale* str. Florida, *A. centrale* str. Israel) and *W. endosymbiont* of *D. melanogaster* (out group) was reconstructed on the basis of concatenated nucleic acid alignment of proteins shared by all species (core genomes) with 100 bootstrap resamplings. The following are represented for each bacterium: genome size (red), number of ORFs (blue), number of predicted T4 effectors (green), number of unique predicted effectors (yellow) and known major hosts (black symbols).

### Comparison of predicted type IV effector repertoires

The similarity of pT4E repertoires was calculated as the mean of (1) the fraction of orthologous pT4Es shared by species A and B, out of all effectors represented in species A, and (2) the fraction of orthologous pT4Es shared by species A and B, out of all effectors represented in species B. Results of side-by-side comparisons were plotted on colour-coded heat maps and sorted according to the order defined by the phylogenetic tree.

### Analysis of *Ehrlichia* spp. genomic architectures and distribution of predicted effectomes

The distribution of pT4Es was analysed in two different ways. First, S4TE 2.0 enables analysis of the predicted effectome distribution according to local gene density (Fig. 5). Second, the distribution predicted effectomes was determined according to their ΔGC content and genome architecture (Fig. 6).

To visualise the distance between each gene and its closest neighbours on the five prime and three prime borders in a single representation, S4TE 2.0 sorted genes into two-dimensional bins defined by the length of their 5’ and 3’ intergenic regions (5’FIR and 3’FIR) (Meyer et al., 2013; Noroy et al., 2019). The colour-coded heat map depicts the gene density distribution. S4TE2.0 used the median length of FIRs to distinguish between gene-dense regions (GDRs) and gene-sparse regions (GSRs) and in-between regions (IBRs). Effectors were then drawn on this heat map according to their flanking intergenic region 5 ‘and 3’. This method makes it possible to visualise the position of pT4Es according to genome density (Fig. 5, Fig S3). An additional Circos graph was used to visualise the distribution of effectors along the genome according to local gene density.

To visualise the local GC content according to the genome architecture in a single representation, we calculated the G+C content of each gene and then subtracted the mean of G+C content of all the genes in the genome, giving the ΔGC content of each gene. The genes were sorted into two-dimensional bins defined by the length of their 5’ and 3’ FIRs. The mean ΔGC content of genes in the same bin was calculated and is represented by a colour-coded heat map. GDRs, GSRs and IBRs were defined as described above. Similarly, pT4ES were plotted on the heat map according to their 5’FIR and 3’FIR. This method makes it possible to visualise the position of pT4Es according to genome architecture and in relation with their GC content (Fig. 6, Fig S4). An additional density graph was used to quantify the ΔGC content of effectors compared to that of other genes. This density graph was constructed using R graphics. The red line represents the density of effectors according to ΔGC content and the black line represents density of all other genes.

### Identification of protein domains in *Ehrlichia* spp

Protein domains of *Ehrlichia* pT4Es were identified using S4TE2.0 and the Pfam database. S4TE2.0 proposes six different modules to find several domains (eukaryotic-like domains, the DUF domain, EPIYA motifs, NLS domains, MLS domains, the prenylation domain, coiled-coil domains). In this study, only EPIYA, NLS, MLS, prenylation and coiled-coil domains were used. We also used PfamScan to search for protein domains in the Pfam database.

We analysed the protein architecture of pT4Es using hive plots, which make it possible to obtain a linear layout of the network of the various domain combinations among these proteins (Krzywinski et al., 2012). The network of protein architectures connected by shared domains was families with Jhive plot software (Krzywinski et al., 2012). Domains were plotted on axes (a1, a2, a3) according to the following rules: On the a1 axis, the number of links between different domains is strictly less than 15; on axis a2, the number of links is between 15 and 40; and on axis a3, the number of links is strictly more than 40 (Fig. 7). To get clearer view of less frequently represented domains, a second hive plot was created by omitting the four most frequently represented domains (NLS, Coiled-coils, EPIYA, Eblock) with the following parameters. On axis a1, the number of links between domains is strictly less than 3; on the a2 axis, the number of links is between 3 and 5; and on the a3 axis, the number of links is strictly higher than 5 (Fig. 8). On both hive plots, the position of the domains on the axes (nodes) is sorted according to the increasing number of links from the centre to the outside.

The analysis of these two graphs prompted us to choose two families of putative effectors for a detailed study of their architectural diversity. The ankyrin-containing putative effector family was chosen to represent one of the two known effectors to be secreted in a type IV dependent manner in *Anaplasmataceae*, i.e. AnkA, ECH_0684 (Fig. 9A). The HATPase_c-containing putative effector family was chosen to represent an effector family with a low number of links and containing a species-specific pT4E (Fig. 10). In order to determine the evolution of Ank-containing predicted T4Es, a time tree (Fig. 9B) was constructed using the Reltime-ML method (Tamura et al., 2012) and the Tamura-Nei model. The analysis involved 11 nucleotide sequences. All positions containing gaps and missing data were eliminated. A total of 1 836 positions comprised the final dataset. Evolutionary analyses were conducted in MEGA7 (Kumar et al., 2016). To check the evolution of homologous gene sequences to AnkA, dot plots were built using dotmatcher software in the EMBOSS package (Rice et al., 2000) with a window size of 50 and a threshold of 50. All positions from the first input sequence were compared with all positions from the second input sequence using a specified substitution matrix. Only the dot-plot comparing ECH-0684 (*x*-axis) and ERGA_CDS_03830 (*y*-axis) sequences is presented here.

## Results

### The host spectrum matches the number of ORFs in *Anaplasmataceae*

The *Anaplasmataceae* phylogenetic tree shows a clear divergence between the two distinct *Ehrlichia* and *Anaplasma* clades (Fig. 1): one clade contains the four *Ehrlichia* species including the most widely studied species *E. chaffeensis* and the other contains the three *Anaplasma* species *A. phagocytophilum, A. marginale* and *A. centrale. Wolbachia endosymbiont* of *Drosophila melanogaster* was included as an outgroup. The length of the genomes ranged from 1.50 Mbp in *A. phagocytophilum* to 1.17 Mbp in *E. chaffeensis, E. muris* and *A. marginale* (Fig. 1). The GC content differed considerably between *Ehrlichia* species and *Anaplasma* species. The GC content of *Ehrlichia* species ranged from 27.5% in *E. ruminantium* to 30.1% in *E. canis*. The GC content of *Anaplasma* species was also highly variable, ranging from 41.7% for *A. phagocytophilum* to 50% for *A. central (data not shown)*. The number of predicted Type IV Effectors (pT4Es) was almost constant (8% of the genome) whatever the genome considered and ranged from 44 for *E. chaffeensis* to 70 for *E. ruminantium* (Fig. 1). S4TE 2.0 identified a total set of 579 pT4Es in the *Ehrlichia* and *Anaplasma* genera. Despite this high number, only five pT4Es are shared by all species (core effectome of *Anaplasmataceae*). The phylogenetic tree of core effectome is in agreement with the core genome phylogenetic tree (Fig. S1) and with the 16S tree (not shown). The pathogenic bacteria belonging to the *Anaplasmataceae* family infect a wide range of hosts. We showed that this broader range of hosts is in accordance with the higher number of ORFs in the genomes (Fig. 1). For instance, *E. chaffeensis*, which contains the higher number of ORFs, has the broadest host range. Inversely, *E. ruminantium* with a larger genome but fewer ORFs, has a more limited host range. However, we point out that the number of species-specific effectors appears to be linked to the size of the genome (Fig. 1).

### Analysis of groups of putative effectors and their corresponding *Anaplasmataceae* hosts suggests several host-specific effectors

To identify a possible link between pT4E repertoires and host specificity, we performed pairwise comparison of effector gene repertoires by calculating the fraction of shared effectors, and then clustered species based on the similarities in their effector pools (Fig. 2). The resulting clusters strongly agree with the phylogenetic clades. Although pT4Es repertoires are versatile, they show a certain level of conservation within a genus and between phylogenetically close species. This is especially true for *Ehrlichia* genus as highlighted by the pale yellow cells in the matrix diagram (Fig. 2). In addition, it is interesting to note that *E. muris* has the same similarity value (0.32) as *A. marginale* with *A. phagocytophilum*. The similarity between *E. muris* and *A. phagocytophilum* could result from the fact that both bacteria can infect a common rodent host. However, although *E. ruminantium, A. marginale* and *A. centrale* have the same hosts (ruminants), there is less similarity between their corresponding repertoires (0.18).

**Figure 2.**
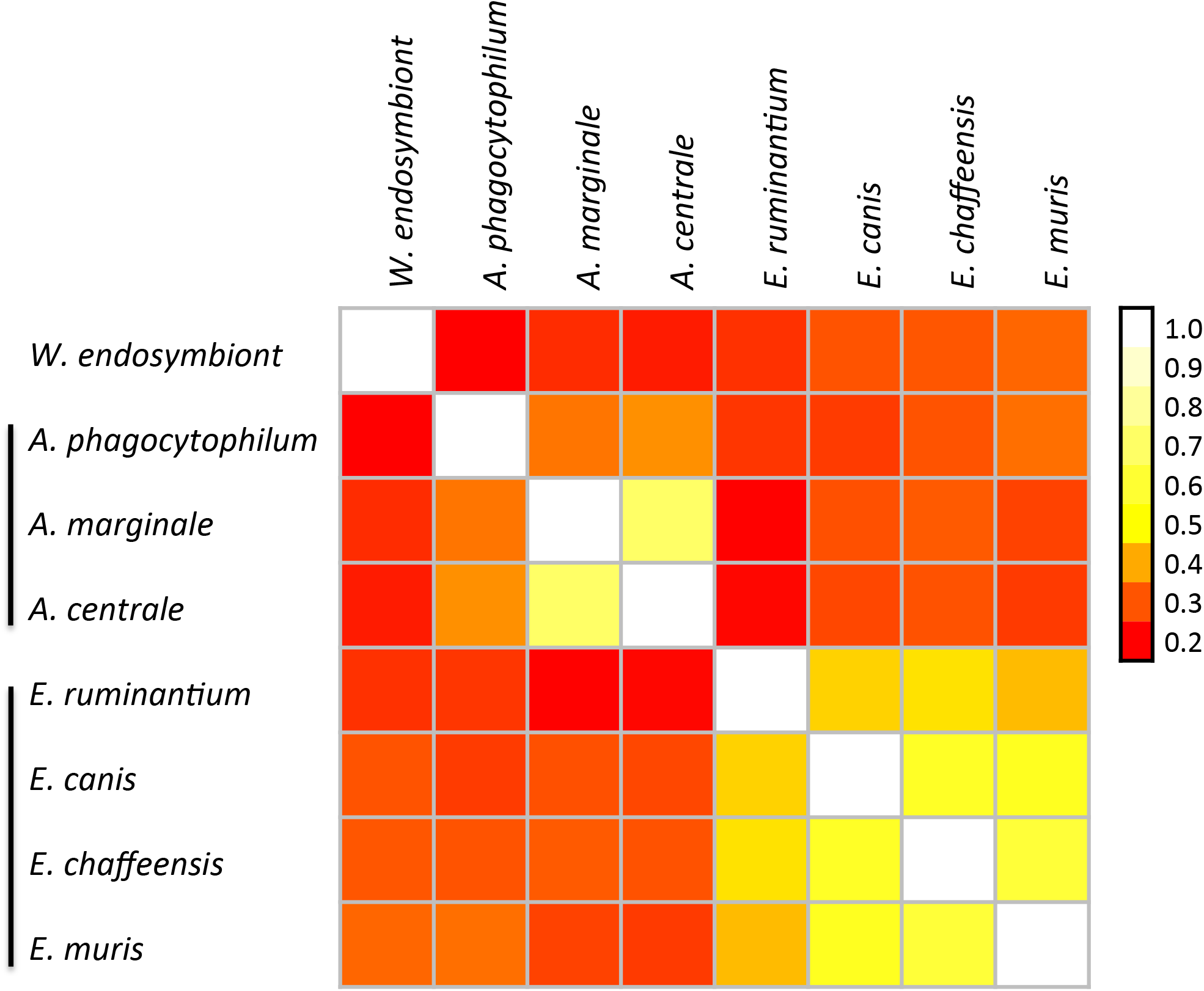
Comparison of the pools of predicted Type IV effectors among *Anaplasmataceae* species revealed strong conservation in *Ehrlichia* spp. The colour gradient represents the similarity between sets of effectors (pale colours mean high similarity). The different species are ordered according to the phylogenetic tree (Fig. 1). Clusters defined on the basis of similar effectors repertoires are marked on the left by black lines.

Within *Ehrlichia* species, *E. chaffeensis* has the broadest host range. Among these preferential hosts, *E. chaffeensis* infects canids and rodents, which are also the specific hosts of *E. canis* and *E. muris* respectively (Fig. 1). Indeed, some preferential hosts of *Ehrlichia* species are shared by *Anaplasma* species. Ruminants are infected by all *Anaplasma* species and by *E. ruminantium*, rodents are infected by *A. phagocytophilum, E. muris* and *E. chaffeensis* and canids are infected by A. *phagocytophilum, E. canis* and *E. chaffeensis* (Fig. 1). The different preferential hosts were plotted on the a2 axes of a hive plot (Fig. 4). The number of species-specific pT4Es of *Ehrlichia* and *Anaplasma* is represented by the size of each node on the a1 and a3 axes, respectively. Links between the a1 or a3 and a2 axes represent effectors possibly involved in host specificity. Links between the a1 and a3 axes represent the homology between *Ehrlichia* species-specific effectors and *Anaplasma* species-specific effectors. For example, *E. ruminantium* (green) shows 33 species-specific pT4Es, which could be involved in host-specificity (ruminants). Thus, among these effectors, some share homology with species-specific pT4Es *of Anaplasma* that are pathogenic on ruminants.

### The *Ehrlichia* pan-genome effector super-repertoire includes core and variable effectors with possibly 30% of species-specific effectors

Figure 3 shows the position of the pT4Es on the genomes of the four *Ehrlichia* species and the homologies between the effectors clearly indicate species-specific effectors (i.e. no orthologues) and effectors belonging to the core effectome (i.e. present in all the strains) (Fig. 3). Interestingly, the colour code highlights an inversion of part of the genomes compared to *E. chaffeensis* as reference (with the biggest number of ORFs). The super repertoire of pT4Es in the pangenome of *Ehrlichia* spp. comprises 52 groups of orthologues, hereafter referred to as effector orthologue groups (EOGs). In other words, 70% of *Ehrlichia* predicted effectors belong to EOGs. Moreover, we found that most predicted effectors were shared by a small subset of species (29 EOGs) and 23 effectors were “core effectors”, i.e. had orthologues in all *Ehrlichia* species (Fig. 3, Fig. S2). Although the evolutionary tree of *Ehrlichia* core effectors (Fig. S1A) is somewhat congruent with the *Ehrlichia* phylogenetic tree (Fig. 1), it is interesting to note some discrepancies in the percentage identity between the 23 EOGs of the core effectome (Fig. S1B). Fourteen EOGs show high sequence identity between *E. canis, E. muris* and *E. ruminantium*. Four EOGs show high pairwise identity between *E. canis* and *E. muris* and *E. chaffeensis* and *E. ruminantium*. A single EOG show strong identity between *E. canis* and *E. muris*. Surprisingly, only four EOGs show strong identity between all pT4Es, particularly between *E. chaffeensis* and *E. muris*. For example, the EOG corresponding to AnkA (ECH_0684, black dot) show high sequence identity between *E. canis, E. muris* and *E. ruminantium* rather than between *E. chaffeensis* and *E. muris* in contrast to what could be inferred from the phylogenetic tree of core effectors (Fig. S1A and B).

**Figure 3.**
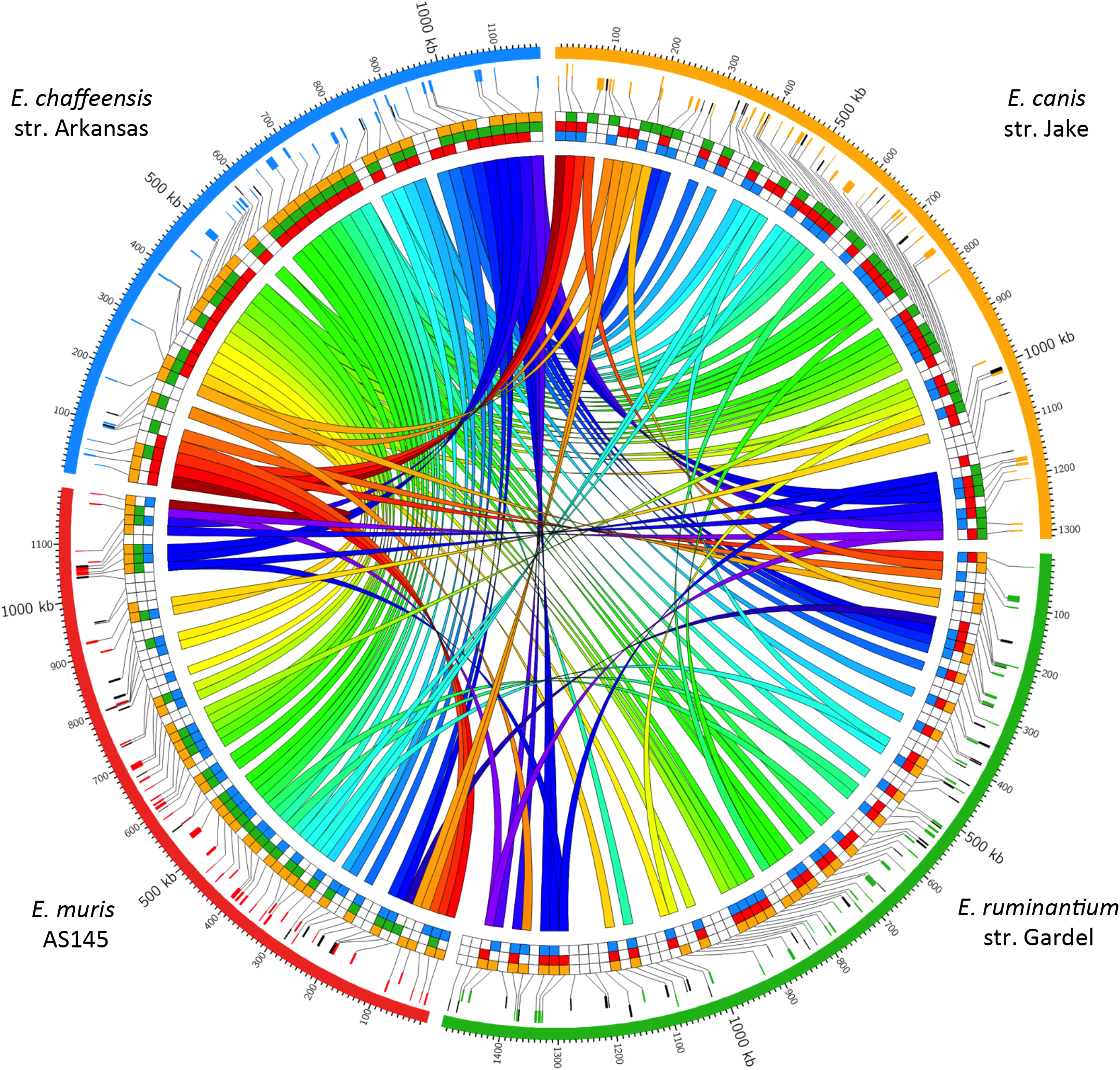
Mapping of *Ehrlichia spp*. predicted Type IV effectors (pT4Es) and their homologies highlights the genomic plasticity of this genus. Genomes of *E. chaffeensis* str. Arkansas (blue), *E. canis* str. Jake (orange), *E. ruminantium* str. Gardel (green) and *E. muris* AS145 (red) are represented in the outer circle of this Circos graph. The second and third circles represent the genes encoding the pT4Es (sense and antisense genes, respectively). The genes are colour coded depending on the genome in which they originated and species-specific genes are in black. Links using a spectral color scheme show homologies between pT4Es of the four genomes. The homologies between the pT4Es of the four genomes are also represented by squares of the corresponding genome colour.

**Figure 4.**
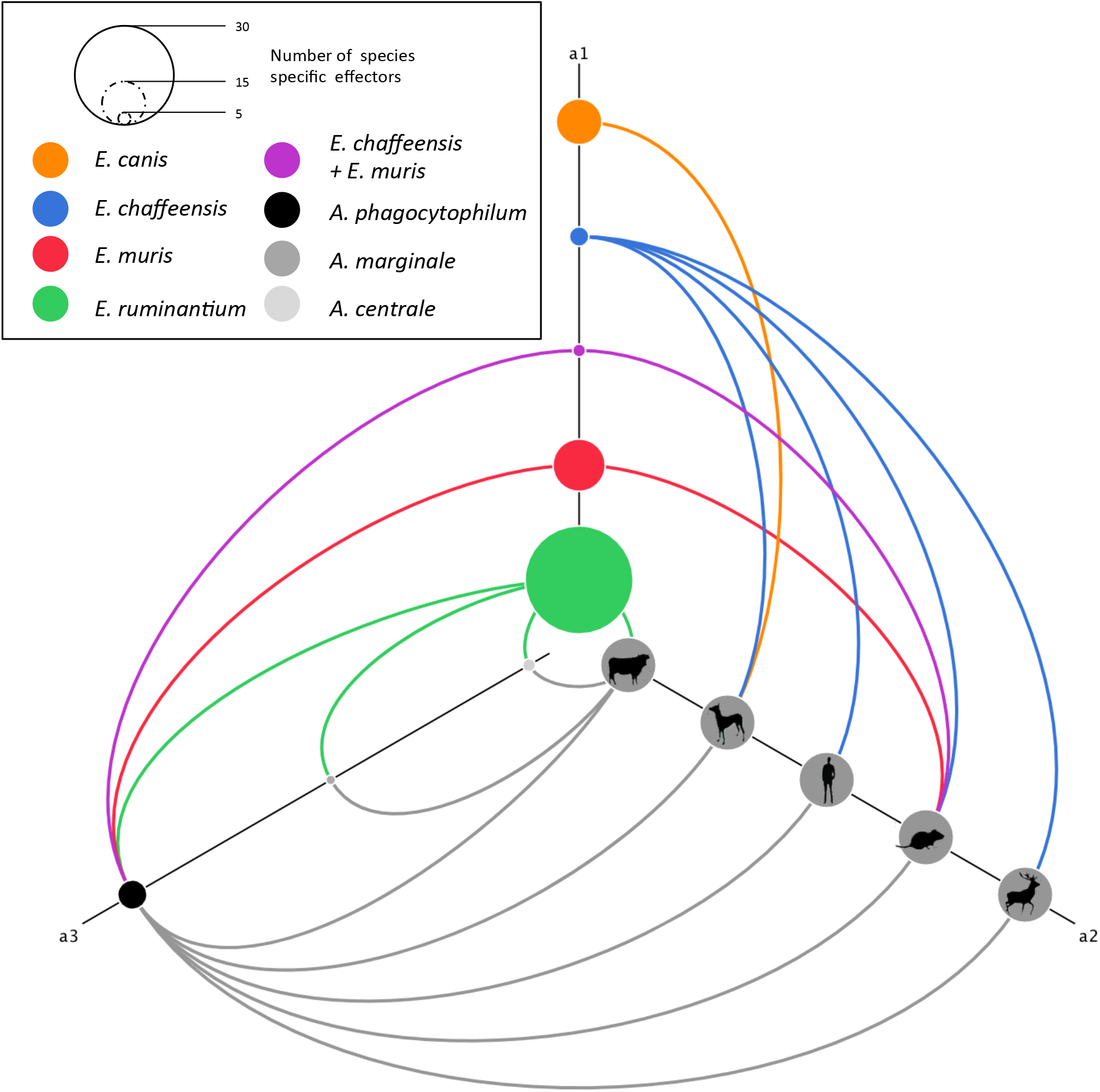
Network analysis of *Anaplasmataceae* species-specific pT4Es and host range suggests the existence of host-specific pT4Es. This network of species-specific pT4Es was drawn using the hive plot algorithm, which is a rational visualization method for drawing networks based on their structural properties. Nodes are mapped to and positioned on radially distributed linear axes and edges are drawn as curved links. *Ehrlichia* species-specific pT4Es are represented by nodes on the a1 axis of the hive plot. The size of each node is linked to the number of species-specific pT4Es for a given *Ehrlichia* species which is colour coded as shown in the upper left rectangle: *E. chaffeensis* str. Arkansas (blue), *E. canis* str. Jake (orange), *E. ruminantium* str. Gardel (green) and *E. muris* AS145 (red). Purple node is the combined subset of pT4Es specific to *E. chaffeensis* str. Arkansas and *E. muris* AS145. *Anaplasma* species-specific pT4E are represented by nodes on the a3 axis whose size and nuance of grey depend on the *Anaplasma* species. The different hosts of these 7 *Anaplasmataceae* bacteria are represented by grey nodes on the a2 axis. Curved links a1-a2 and a3-a2 show the putative host specificity of each bacterium. Links between a1-a3 represent shared host-specific effectors. The colour of each link is related to the node from which it emerges.

The analysis of effectors revealed that 30% of them (69) were only observed in one of the *Ehrlichia* species analysed. The species with the highest number of unique pT4Es is *E. ruminantium*, with 33 species-specific effectors. Notably, each *Ehrlichia* genome contains at least six species-specific effectors (Fig. 3, Fig. S2). In addition, species-specific effectors appear to be randomly located in the genome relative to other effectors. This is notable in the genome of *E. ruminantium*, which has the higher number of species-specific effectors (Fig. 3, Fig. S2).

### Predicted type IV effectors of *Ehrlichia* species are overrepresented in gene sparse regions and in high GC content regions of the genome

In order to understand how genomic plasticity influences the distribution of pT4Es, we first analysed the genome architecture of *Ehrlichia* species by looking at local gene density (Fig. 5, Fig. S3). The gene architecture of *E. canis* shows 29.2% of genes in gene dense regions (GDRs) and in gene sparse regions (GSRs) while 41.6% of genes are in ‘in between’ regions (IBRs) (Fig. 5). This genome architecture is also representative of other *Ehrlichia* species (Fig. S3). Although 29.2% of *E. canis* genes belong to GSRs, 42% of pT4Es and 42.9% of species-specific pT4Es are in GSRs (Fig. 5). Thus, compared to the whole genome, the GSR showed a 1.44-fold enrichment in candidate type IV effector genes. The proportion of candidate T4Es in IBRs is not significantly different, with 42.8% of *E. canis* genes belonging to IBRs and 43.5% of pT4Es - 42.9% of species-specific pT4Es - in IBRs (Fig. 5). Consequently, the proportion of candidate T4Es in GDRs is lower than the proportion of genes of the whole genome. These results suggest that plastic regions with low gene density harbour pathogenicity genes and could play a role in host-bacteria interactions.

**Figure 5.**
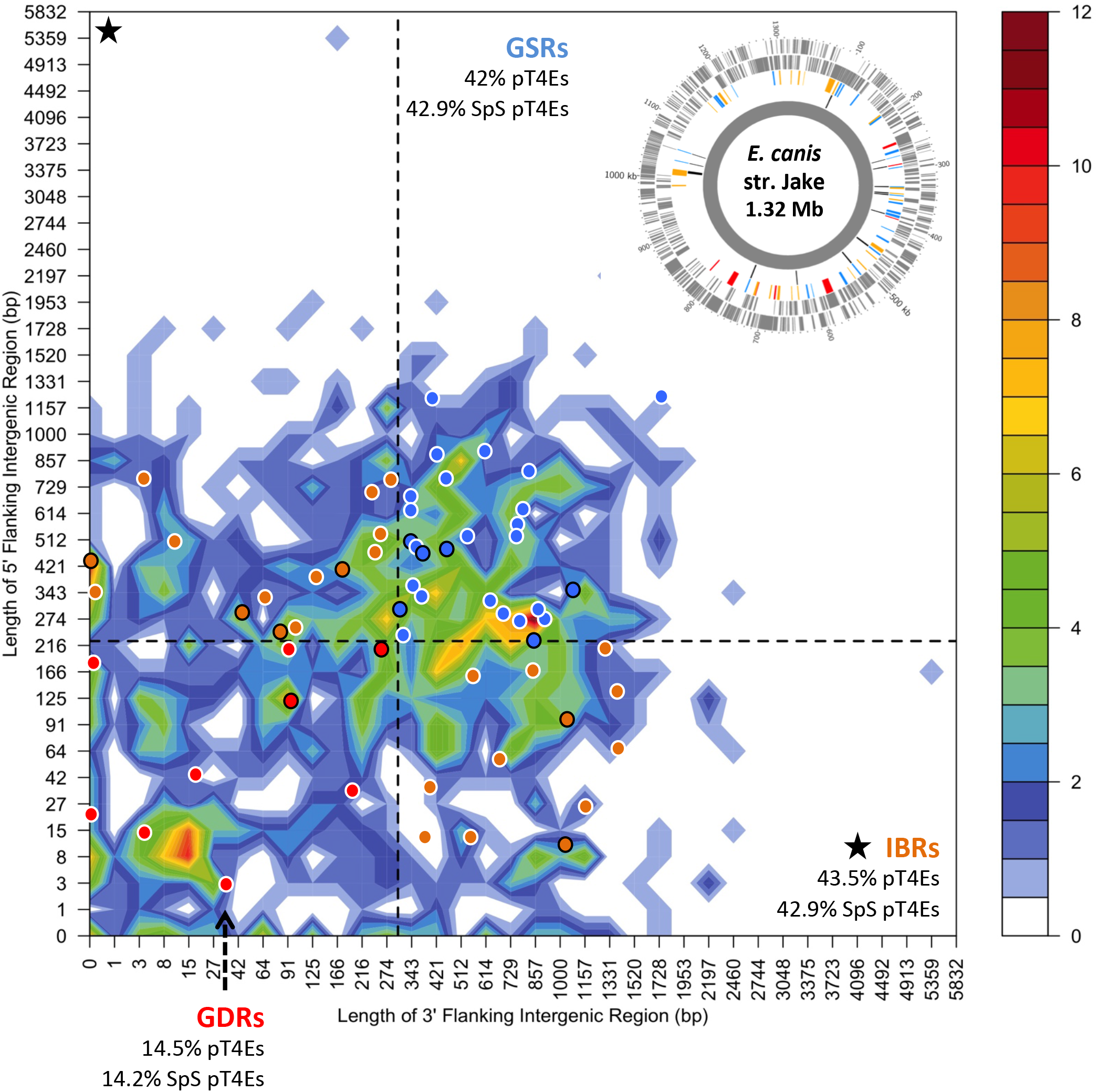
The distribution of *Ehrlichia* type IV effectomes according to local gene density shows an enrichment of pT4Es in gene sparse regions. Distribution of *E. canis str*. Jake genes according to the length of their flanking intergenic regions (FIRs). All *E. canis* genes were sorted in two-dimensional bins according to the length of their 5⍰ (*y*-axis) and 3⍰ (*x*-axis) FIRs. The number of genes in the bins is represented by a colour-coded density graph. Genes whose FIRs were both longer than the median length of FIRs were considered as gene-sparse region (GSR) genes. Genes whose FIRs were both below the median value were considered as gene-dense region (GDR) genes. In between region (IBR) genes are genes with a long 5⍰ FIR and short 3⍰ FIR, and inversely. For *E. canis*, this median value is 225 bp for 5⍰ FIRs and 304 bp for 3⍰ FIRs. The dashed line showing the median length of FIR delimits the genes in GSR, GDR and IBR. Candidate effectors predicted using the S4TE 2.0 algorithm were *s* plotted on this distribution according to their own 3⍰ and 5⍰ FIRs. A colour was assigned to each of the three following groups: red to GDRs, orange to IBRs, and blue to GSRs. Specific pT4Es are represented by a dot circled in black. In the top right corner, a Circos graph shows the distribution of *E. canis* str. Jake putative effectors along the genome. The outermost and second circles (in grey) represent *E. canis* antisense and sense genes, respectively. The third and innermost circles represent pT4Es. The black, red, orange and blue colour of each putative T4 effector corresponds to species-specific effectors located in GDRs, IBRs and GSRs, respectively.

Contemporary methods used to infer horizontal gene transfer events are based on analyses of genomic sequence data. One interesting method consists in searching for a section of a genome that significantly differs from the genomic average, such as GC content (Lawrence, 2002). To understand how genomic plasticity and horizontal gene transfers could influence the distribution of predicted T4Es, we analysed the genome architecture *of Ehrlichia* species according to the local GC content (Fig. 6, Fig. S4). These figures show that there is an enrichment of pT4Es in regions with high GC content. Indeed, 67.7%, 69.4% and 73.3% of the pT4Es of *E. canis, E. muris* and *E. chaffeensis*, respectively, were found to have a higher ΔGC than the average of the other genes (50% of genes less than or equal to the average ΔGC). It is noteworthy that species-specific pT4Es are also over-represented in high GC content regions with 53.3% and 75% for *E. canis* and *E. muris*, respectively. For *E. ruminantium*, the difference in GC content between effectors and other genes is less pronounced because this genome has a mosaic structure with zones of high density GC content (in red in Fig. S4A). In addition, unlike the curve of the other genes in the genome (normal curve), several density peaks are highlighted by the ΔGC-density curve of putative effectors (Fig. 6 Fig. S4). These results show that pT4Es (and species-specific pT4Es) are mostly located in high GC content regions that possibly favoured the acquisition of new genes during the evolution of the host-pathogen relationship.

**Figure 6.**
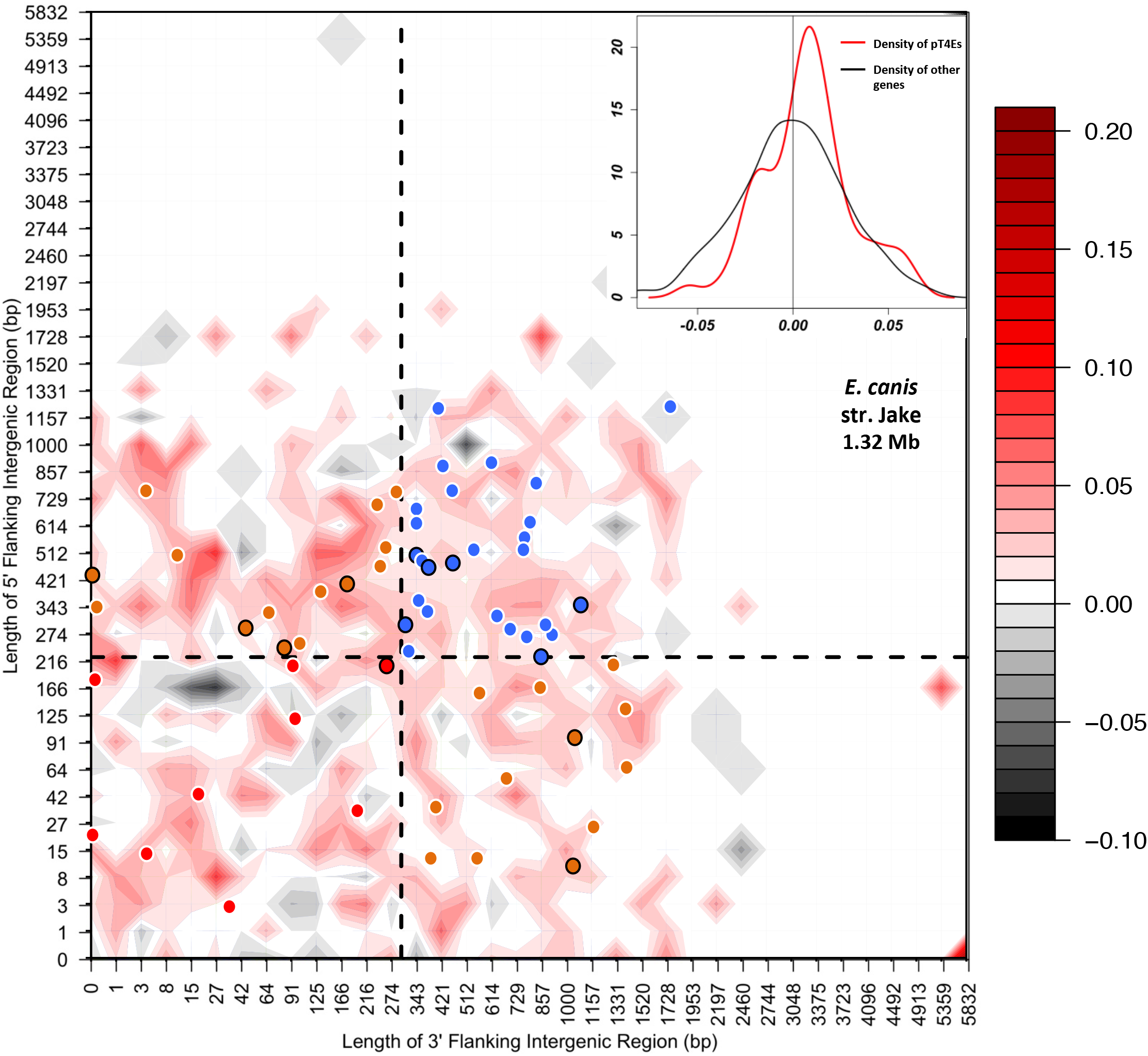
The distribution of *Ehrlichia* type IV effectomes according to local gene density and ΔGC content shows an enrichment of pT4Es in high GC content regions. Distribution of *E. canis* str. Jake genes according to the length of their flanking intergenic regions (FIRs). All *E. canis* genes were sorted in two-dimensional bins according to the length of their 5⍰ (*y*-axis) and 3⍰ (*x*-axis) FIR lengths. For each gene, the ΔGC content was calculated by subtracting the GC content of a gene by the average of GC content of all the genes. The mean of ΔGC of genes in the bins is represented by a colour-coded density graph. GSR, GDR and IBR were defined as described for the analysis of local gene density. A colour was assigned to each of the three following groups: red to GDRs, orange to IBRs, and blue to GSRs. Specific pT4Es are represented with a dot circled in black. In the top right corner, a density graph indicates the density of pT4Es according to ΔGC content (red line) and the density of other genes (black line).

### Domain shuffling seems to play a major role in effector evolution

Previous studies of T4Es revealed that they harbour numerous eukaryotic domains as well as effector-specific domains (Burstein et al., 2016; Gomez-Valero et al., 2011). The high number of effectors predicted by S4TE 2.0 made it possible to identify and analyse conserved effector domains across the *Ehrlichia* genus. We identified the domains present in *Ehrlichia* pT4Es using a similarity search of Pfam databases, and outputs of S4TE 2.0 software. Conserved domains were detected in 97% of the pT4Es. A total of 116 distinct domains were identified by the two methods combined. When analysing the protein architectures (different domain combinations), we noticed that the same domains were often shared among different architectures. We visualised this phenomenon as a network of protein architectures connected by shared domains (Fig. 7). The network of protein architecture of pT4Es was drawn using the hive plot algorithm. The network clearly demonstrates that several domains are present in numerous effectors (indicated by the occurrence and the width of the links), as well as in numerous different architectures (indicated by the number of links). The most common domains in *Ehrlichia* T4Es are domains for protein subcellular location like NLS or MLS, domains using protein-protein interactions like coiled-coils or Ank domains, and domains for post-translational modification like EPIYA or HATPase_c. NLS, which are known to address proteins to the nucleus of eukaryotes, were found in 67% (162) of the pT4Es. The coiled-coils domains, known to mediate protein-protein interactions, were found in 36% (87) of pT4Es. The EPIYA domains, known to be phosphorylation sites, were found in 31% (76) of pT4Es. Finally, the Eblock domain, known as a secretion signal in *Legionella pneumophila*, was found in 29% (70) of pT4Es. In addition to these four most abundant and best-known domains, the network represents a wealth of domains with both known and unknown functions. To facilitate the reading of the least represented domains, a second network was built by removing the four most frequently represented domains described before (NLS, coiled-coils, EPIYA and Eblock) (Fig. 8). Interestingly, some domains like Ank repeats disappeared from the network. In other words, Ank repeat domains were always associated with at least one of the four most widely represented domains.

**Figure 7.**
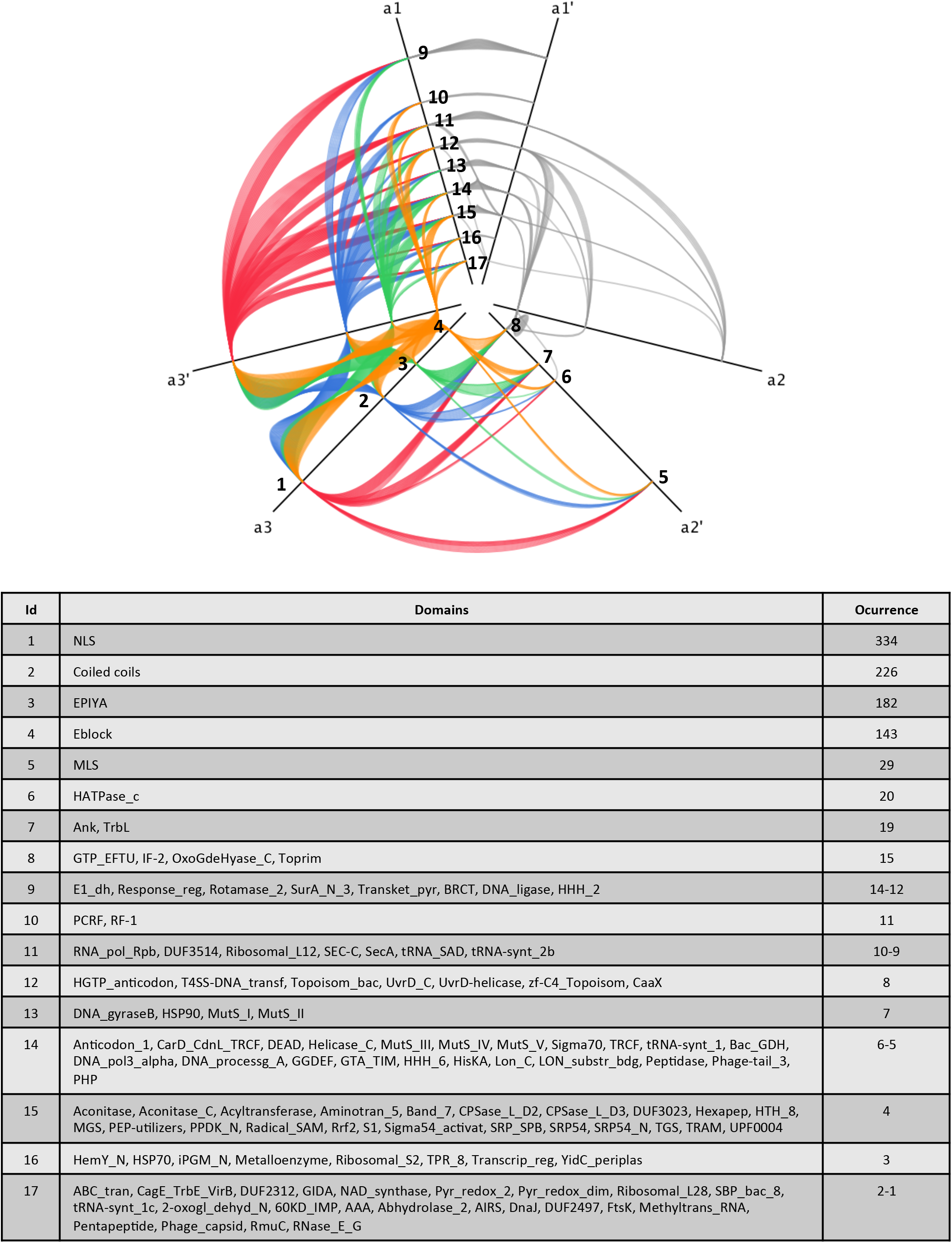
Protein architecture network of *Ehrlichia* pT4Es shows a large number of interactions between protein domains. This network of protein architecture of pT4Es is drawn using the hive plot algorithm, which is a rational visualization method for drawing networks based on their structural properties. Nodes are mapped to and positioned on radially distributed linear axes, and edges are drawn as curved links. Each node represents a specific domain or a list of specific domains (see table) found in *Ehrlichia* T4Es predicted by S4TE 2.0. Links between domains represent the association of these domains in the architecture of *Ehrlichia* pT4Es. Links between NLS, Coiled-coils, EPIYA and Eblock domains (the most abundant protein domains in *Ehrlichia*) and other nodes are red, blue, green and orange, respectively. Other links are pale grey. The table of domains identifies the protein domains for each node (numbered) and their occurrences in *Ehrlichia* spp predicted T4 effectomes. All the domains are ranked on the three axes (a1, a2, a3) according to the number of their links. Let X be the number of links between one domain and the others, X < 15 was represented on a1, 15 ≥ X ≤ 40 was represented on a2, and X > 40 was represented on a3, thus defining the most abundant domains, a1’, a2’ and a3’ split axes were drawn to represent the links between domains located on the same axis.

**Figure 8.**
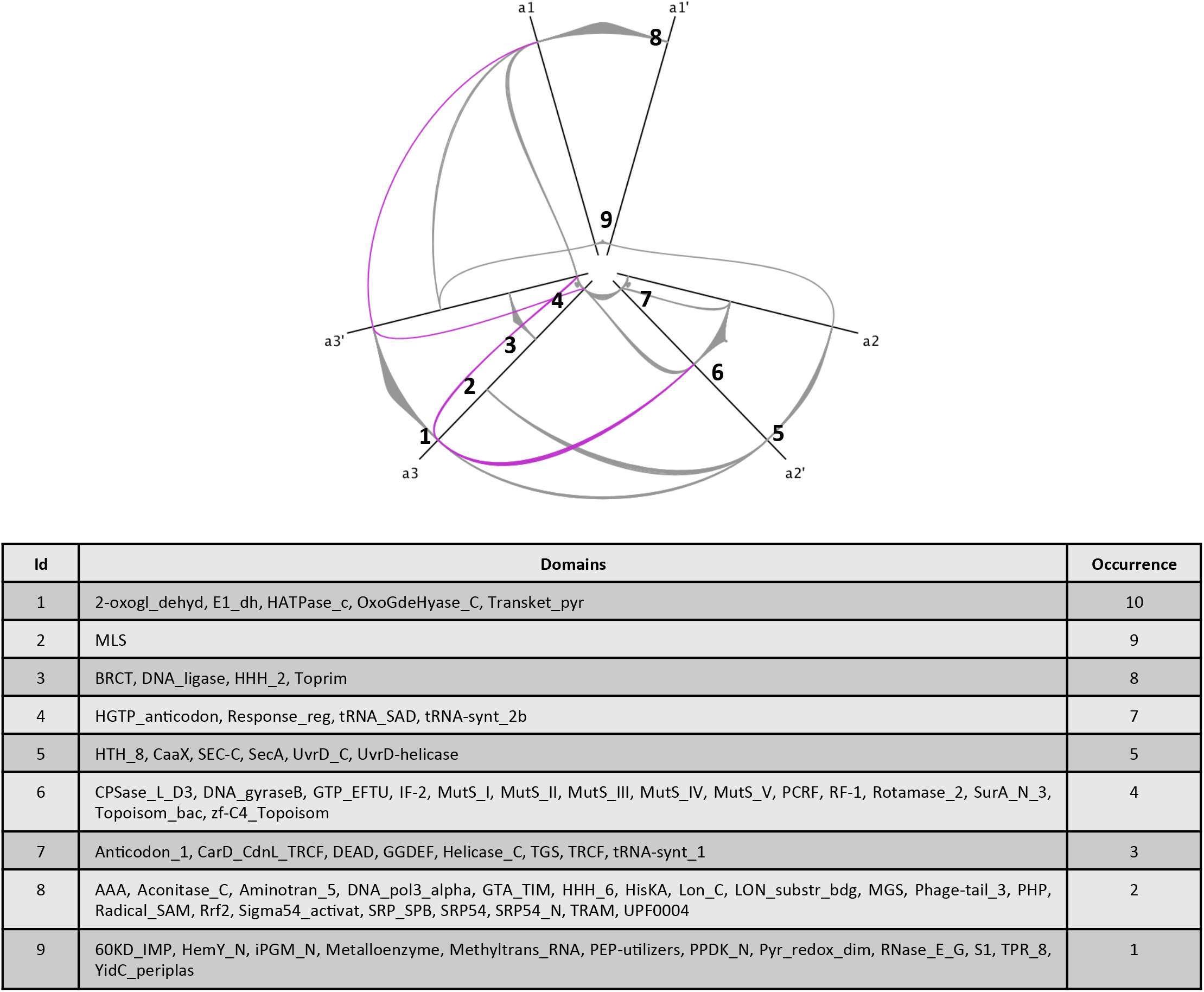
Protein architecture network of putative effectors for rarely occurring domains. This network of pT4E protein architectures was drawn using the hive plot algorithm to produce a rational visualization method based only on the network structural properties. Each node represents a specific domain or a list of specific domains (see table) found in *Ehrlichia* T4Es predicted by S4TE 2.0. Links between domains represents the association between these domains in the architecture *of Ehrlichia* pT4Es. Nodes representing the most abundant domains presented in fig. 7 (NLS, Coiled-coils, EPIYA and Eblock domains) and their corresponding links to other nodes are not included in this graph to highlight the less abundant protein domains in *Ehrlichia* spp predicted T4 effectomes. The table of domains identifies the protein domains for each node (numbered) and their occurrences. All the domains are ranked on the three axes (a1, a2, a3) according to the number of their links. Let X be the number of links between one domain and the others, X < 3 was represented on axis a1, 3 ≥ X ≤ 5 was represented on axis a2, and X > 5 was represented on axis a3.

In *Ehrlichia* species, Ankyrin repeats (Ank) appear to be associated with two different domains (NLS and EPIYA). Overall Anks were found in 11 pT4Es in the four species (Fig. 9A). Although there is strong homology between different Ank-containing T4Es, differences in the position of the different protein domains were observed. For example, the first of Ank-containing pT4Es, Ecaj_0365, EMUR_01925 and ERGA_CDS_03830 are homologous with the known effector AnkA ECH_0684 of *E. chaffeensis*, but the position of the Ank, EPIYA and NLS domains differs, some domains are even absent in certain proteins (Fig. 9A). Further, the analysis of ECH_0684 and ERGA_CDS_03830 revealed seven duplications of about 200 amino acids of ECH_0684 (from 200 to 400) in ERGA_CDS_03830 (showed by the seven groups of diagonals on the upper side of the main diagonal in Fig. 9C). Despite these differences, we were able to highlight three groups of proteins with the time tree (Fig. 9B). The second group on Ank-containing pT4Es comprises Ecaj_0221, ECH_0877 and ERGA_CDS_02160 that show conserved architectures, while the last group of Ank-containing pT4Es show versatile architectures (Fig. 9A). The evolutionary tree of the Ank-containing pT4Es is congruent with the phylogenetic tree of *Anaplasmataceae* species (Fig. 1) for the first (AnkA orthologues) and second groups. Moreover, these two groups appeared to derive from a common ancestral gene and their difference may be linked to different coevolution in the bacterium and its host.

**Figure 9.**
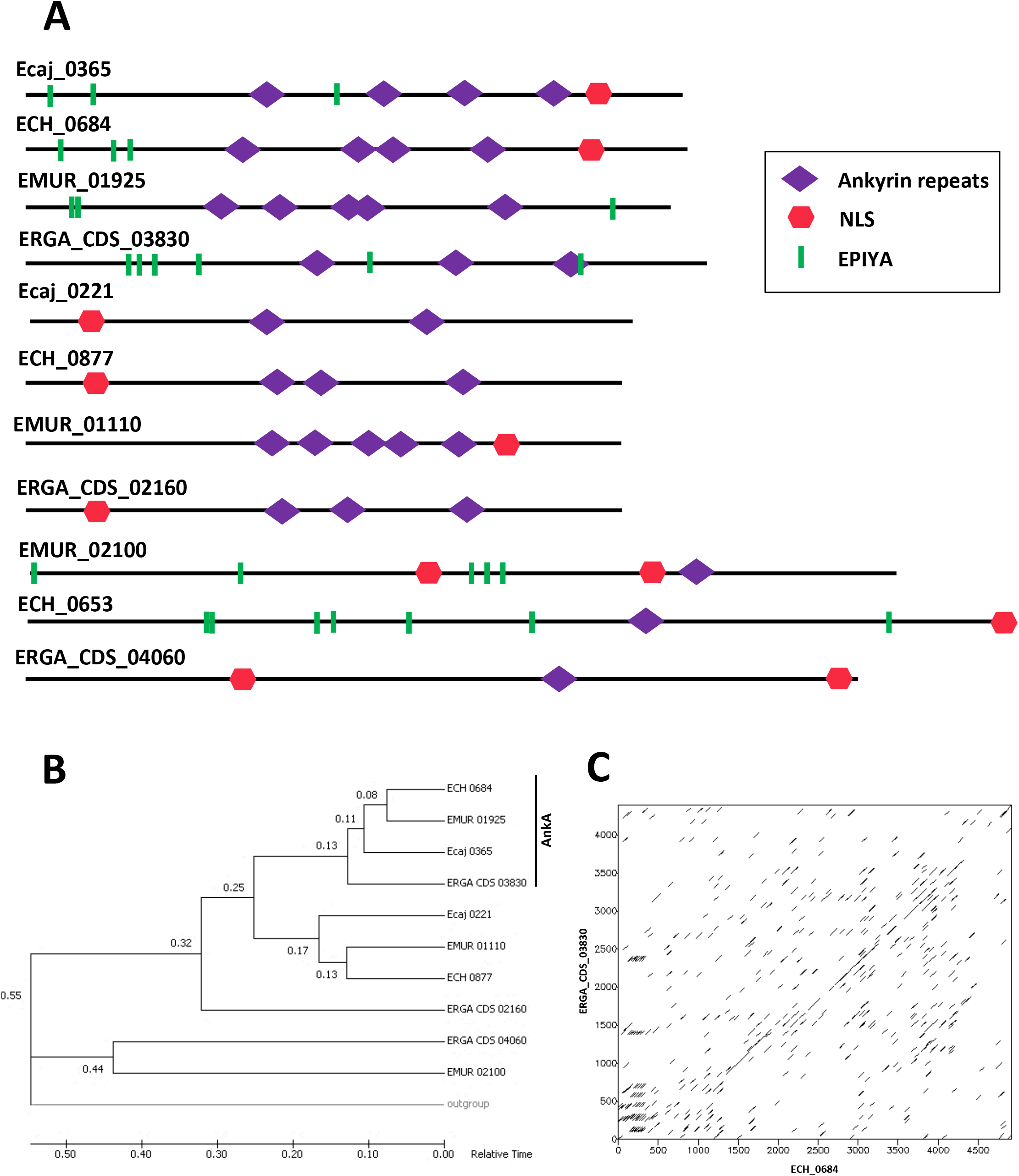
Ankyrin-containing predicted type IV effectors show diverse architectures and inter- and intragenic rearrangements in *Ehrlichia* spp. Each protein in *Ehrlichia* spp. pT4E whose architecture includes an ankyrin domain is represented. **B.** Relative time phylogenetic tree build from 11 nucleotide sequences of Ankyrin-containing predicted type IV effectors (pT4Es). ECH_0653 was used as outgroup. The numbers in front of each node represent the relative theoretical time from the putative common ancestor of two branches. Evolutionary analyses were conducted in MEGA7 (Tamura et al., 2012). **C.** Dot plot of regions of similarities between ECH-0684 (*x*-axis) and ERGA_CDS_03830 (*y*-axis). This graph was constructed with the dotmatcher software included in the EMBOSS package, where all positions from the first input sequence are compared with all positions from the second input sequence using a specified substitution matrix and using a window size of 50 and a threshold of 50. The two sequences are the axes of the rectangular dotplot. Wherever there is “similarity” between a position from each sequence a dot is plotted.

Another family of special interest is *Ehrlichia* pT4Es whose architecture harbours a HATPase_c domain. The HATPase_c was found in only six pT4Es but is present in three different architectures. Two domain architectures are conserved among the different species of *Ehrlichia*, while the third domain architecture is only represented by *E. ruminantium* ERGA_CDS_03390, a species-specific pT4E (Fig. 10). Some domains, including HisKA, HSP90, Toprim, DNA_J, Reponse_ reg, were found adjacent to the HATPase_c domain. HATPase_c is present in several ATP-binding proteins including histidine kinase, DNA gyrase B or the heat shock protein HSP 90. The variety of protein domains carried by these pT4Es indicates that this family may have a different mode of action. It is interesting to note that effectors with domains related to DNA binding (DNA_j, Toprim) also have one or more NLS domain (Fig. 10A).

**Figure 10.**
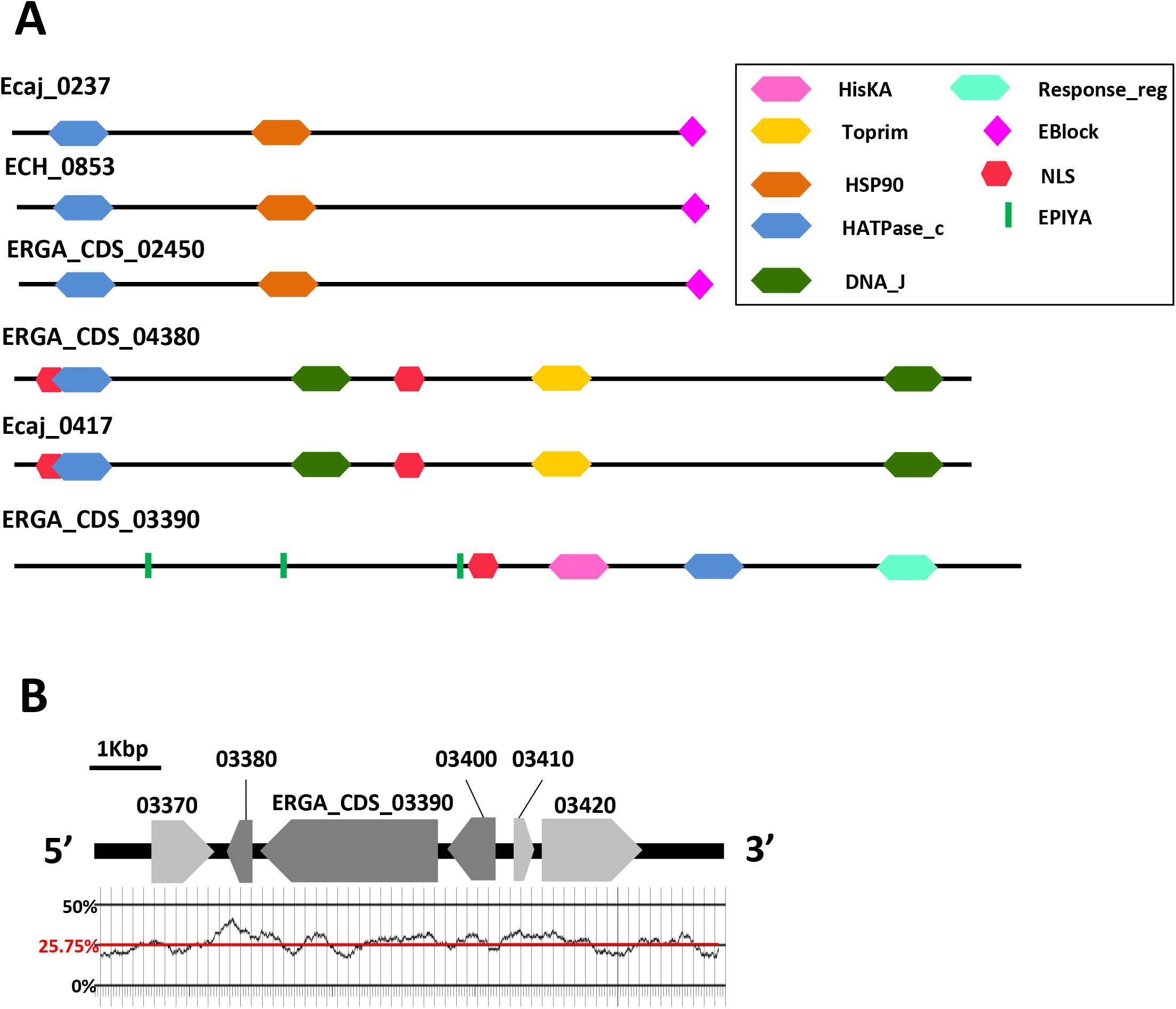
The strong domain diversity of HATPase_c-containing putative type IV effectors defines three conserved families of effectors in *Ehrlichia* spp. **A.** Each protein in *Ehrlichia* spp. type IV effectome whose architecture includes a HATPase_c domain is represented. B. Representation of the genomic context surrounding pT4E ERGA_CDS_03390 between position 557733 and 563669 of *E. ruminantium* str. Gardel genome. Pale grey arrows represent sense genes and dark grey arrows represent anti-sense genes. The GC content of this region of the genome was calculated using 200 bp windows and is represented by the black curve. The average GC content of this gene cluster (25.75 % of GC) is indicated by the horizontal red line.

When we analysed the genomic environment of the ‘orphan’ pT4E ERGA_CDS_03390 in more depth, we found that this gene is part of a cluster of six genes, three of which are sense and three (including ERGA_CDS_03390) are antisense (Fig. 10B). For 5’ to 3’ the cluster is composed as follows (i) a sense gene (ERGA_CDS_03370) encoding an enzyme involved in the methylerythritol phosphate pathway (isopentenyl-pyrophosphate biosynthetic metabolic pathway), (ii) an antisense gene (ERGA_CDS_03380) encoding CutA, an ion transporter, (iii) ERGA_CDS_03390 encoding a protein homologous to VirA, which is a protein sensor secreted by *Agrobacterium tumefaciens*, (iv) the last antisense gene (ERGA_CDS_03400) coding for a putative O-methyltransferase, (v) a sense gene encoding a hypothetical protein, and (vi) a sense gene encoding a NAD(P)H-hydratase dehydratase, which acts on both hydrated NADH and hydrated NADPH (Fig. 10B). Although the average GC content of this part of genome is 25.75%, the antisense cluster showed several peaks of GC content (mean GC content of 26.4% for the three antisense genes).

## Discussion

The obligatory intracellular pathogens of the *Ehrlichia* genus cause fatal infectious diseases. Although they are strictly restricted to their mammalian host or tick vector cells, *Ehrlichia* spp. exhibit marked differences in host range (Moumene and Meyer 2016). Moreover, despite the reductive evolution of their genome, they all have evolved a versatile type IV secretion system which translocates a repertoire of effector proteins (T4Es) into the cytoplasm of host cells, hijacking innate immunity, and manipulating numerous cellular pathways to their own advantage (Rikihisa, 2017). Thus, it was tempting to speculate that T4E repertoires shape the *Ehrlichia* spp. host range. In order to tentatively identify candidate genes involved in host specificity, we predicted *Ehrlichia* spp. T4Es and analysed these T4E gene repertoires.

Here, we report that the number of hosts infected by the bacteria appears to be related to the number of ORFs in their genome. Bacteria can acquire and maintain a diverse repertoire of accessory (variable) genes as a key feature to better adapt to changing environments and to colonise a wider range of ecological niches. This is a well-known phenomenon among opportunistic or ubiquitous pathogens which have bigger genomes than bacteria living in specific more constrained specific environments (Bobay and Ochman, 2017). However, for the broad host spectrum *E. chaffeensis, we* only found six variable pT4Es that are species-specific effectors. The host range thus does not appear to be only related to effector repertoires but to a wider set of genes, and may also be governed by other pathogenicity factors such as host cell adhesion or other type of effectors which ensure bacterial survival (Luo et al., 2011; Farris et al., 2017). Similar observations have been made when *Pseudomonas syringae* pathovars T3E repertoires were compared (Baltrus et al., 2012), thereby reinforcing the hypothesis that a complex genetic basis underlies host range evolution in bacterial pathogens. Some of the variable effectors *of the Ehrlichia* pangenome effector super-repertoire, species-specific effectors, could harbour the strongest virulence phenotypes. Although we identified species-specific genes but not strong host-specificity candidates, we highlighted a strong similarity between the effector repertoires of *E. muris* and *A. phagocytophilum*, two bacteria that share the same rodent host. Similarly, we found 12 *Ehrlichia*-specific effectors in *Anaplasma* that share the same hosts, thus supporting the hypothesis of a coevolution model governed by molecular dialogues between effector repertoires and host immunity. Regarding the question of effector adaptation to different hosts, most *Ehrlichia* effectors may function as ‘generalists’ in a broad range of hosts, and immune response to effectors may be the primary driver of effector repertoire diversification (Tago and Meyer, 2016).

On the other hand, the function of the core effectors remains largely unknown but their high conservation within the evolution of the *Ehrlichia* family shows that the core effectome may fulfil a critical function during infection. We further reveal that T4E gene repertoires of these pathogens comprise core and variable gene suites, which probably have distinct roles in pathogenicity and different evolutionary histories. A frequent hallmark of genes with an extrinsic origin is the difference in GC content of these genes compared with the mean content of the host genome (Dufraigne et al., 2005; Kado, 2009). Around 70% of T4E genes exhibit a high mean GC%, whereas the genomic mean content in *Ehrlichia* spp. is around 28%. The GC content of *Ehrlichia* predicted T4Es is consistently higher than the genomic GC content, suggesting these genes were recently acquired from an exogenous source by horizontal gene transfer (HGT), possibly from natural hosts or the natural tick vector of *Ehrlichia*, which are typically characterised by high GC content. In addition, in the case of the 69 *Ehrlichia* species-specific effectors, the fact that none showed significant sequence similarity to another *Ehrlichia-encoded* protein underlines the magnitude of the functional novelty of the putative effectors we found. The high GC content of species-specific effectors combined with the fact that most of them contain a secretion signal (Eblock), suggest that recently acquired genes can adapt to function as effectors in a relatively short evolutionary time and could be linked to jumps of the host barrier. Our results also suggest that a majority of species-specific effectors may be part of the flexible genome in *Ehrlichia* spp.

Despite their obligate intracellular nature, *Ehrlichia* ssp. show a high level of genomic plasticity. As an example, the large numbers of species-specific pT4Es of *E. ruminantium* appear to be randomly inserted into the genome and may therefore be the result of multiple acquisition events. Similarly, we observed a large chromosomal inversion in the genome of *E. chaffeensis*. It has been shown in numerous bacteria that effector-coding genes in close genomic proximity can function together in the host cell (Ingmundson et al., 2007; Kubori et al., 2010; Siamer and Dehio, 2015). Our search for pairs of effectors in *Ehrlichia* spp. led to the identification of 18 pairs of which only seven were present in at least one other *Ehrlichia* genome (data not shown). This suggests a certain degree of co-evolution of these pairs during the evolution of *Ehrlichia* genus, indicating they could have related functions important for *Ehrlichia* spp. pathogenesis. Interestingly, one of these pairs was found in three *Ehrlichia* genomes, and one of the gene had orthologues in the four *Ehrlichia* species and encoded a lipoprotein which plays a role in *Ehrlichia* pathogenesis (Huang et al., 2008). While analysing the protein architectures (different domain combinations), we noticed that the same domains were often shared by different architectures. We visualised this phenomenon using hive plots to reveal the network of protein architectures based on their structural properties. This network clearly shows that several domains are present in many effectors and form different architectures. Among the most connected architectures, we found well-known effector domains including NLS domains, coiled-coil domains, EPIYA domains and E-block domains. Similarly, among the less abundant architectures, we found domains including HATPase_c domains, MLS domains, and BRCT domains, which are closely connected. These networks represent a plethora of domains of known and unknown functions, which could provide information on possible effector functions.

Ankyrin-repeat (Ank) proteins mediate protein-protein interactions involved in a multitude of host processes. Ank domain-containing effectors are crucial for the pathogenesis of several obligate intracellular bacteria (Rikihisa and Lin, 2010). In addition, when associated with NLS domains, such as *Anaplasma phagocytophilum* AnkA, some Ank-containing effectors target the host cell nucleus to directly interfere with the host defences at the gene and chromatin level (Bierne and Cossart, 2012). In *Ehrlichia* spp., we showed that the Ank-containing pT4Es are associated with few other domains (NLS and EPIYA) but they show a myriad of different protein architectures with some sequence rearrangements (duplications). Hence, the sequence and position of the different domains on the Ank-containing effectors point to the many protein interactions and cellular functions they may have in the host cell. Thus, the modular architecture of Ank-containing effectors, as well as the intragenic recombination we showed, may have played a critical role in the evolution of *Ehrlichia* virulence related to host specificity, as shown for SidJ in *Legionella pneumophila* (Costa et al., 2014). Such high modularity may also be linked to some loss of function, as we showed for *E. ruminantium* ERGA_CDS_03830, which is not secreted in a type IV-dependent manner in the *Legionella pneumophila* heterologous system (data not shown). The great variety and polymorphism of Ank-containing pT4Es may be crucial for increasing the *Ehrlichia* spp. genetic pool and may contribute to the resilience of the bacteria. Further experiments should be conducted to clarify the importance of this polymorphism.

The HATPase_c domain is found in several ATP-binding proteins including histidine kinase and DNA gyrase. This domain is of particular interest because it could define a novel effector function. Even though numerous type III effector kinases have been characterised (Dean, 2011), no HATPase_c-containing T4E has been described to date. These effectors could be highly novel proteins with previously unseen biochemical properties. In contrast to Ank-containing effectors, this family of pT4E shows conserved protein architectures with a wide variety of domain associations. ERGA_CDS_03390 is a species-specific pT4Es of the *E. ruminantium* containing the HATPase_c domain. Even if orthologues of this gene are present in *E. chaffeensis* and *E. canis*, they were not predicted as T4Es by SATE 2.0 software. Indeed, ERGA_CDS_03390 had a score of 101 while ECH_0755 and Ecaj_0319 (homologous proteins of ERGA_CDS_03390) scored below 72, the threshold of S4TE 2.0. The differential score of ERGA_CDS_03390 is due to the presence of a PmrA promoter domain upstream from the gene. The response regulator PmrA is a major regulator of the *icm/dotA* type IV secretion system in *Legionella pneumophila* and *Coxiella burnetii* (Zusman et al., 2007). PmrA is important for the regulation of the effectors and could enable positive regulation of the ERGA_CDS_03390 gene. Its absence in *E. chaffeensis* and *E. canis* and the disappearance of the gene in *E. muris* suggests that this gene no longer has any function in these species. In this sense, ERGA_CDS_03390 could encode a putative effector related to host specificity. ERGA_CDS_03390 is homologous to VirA. VirA is a sensory component of a two-component signal transduction system. This protein is composed of a HisKA domain, a HATPase_c domain and a response_reg domain. The response_reg domain receives the signal from the sensor partner in bacterial two-component systems. It is usually found in the N-terminal of a DNA binding effector domain (Pao and Saier, 1995). Upstream and downstream from this gene are two antisense genes: (I) a gene encoding the periplasmic divalent cation tolerance protein CutA (ERGA_CDS_03380) and (ii) an O-methyltransferase (ERGA_CDS_03400). These two genes may be related to the function of VirA. Indeed, in *A. tumefaciens* crosstalk has been demonstrated between chemotaxis and virulence induction signalling thanks to the interaction modulated by CheR methyltransferase between VirA and transmembrane chemoreceptors MCPs (Guo et al., 2017). In *E. ruminantium*, the three-gene cluster containing ERGA_CDS_03390 pT4E could therefore be related to the perception of the external environment and the subsequent induction of virulence. Moreover, this cluster is antisense, surrounded by ERGA_CDS_03370 and ERGA_CDS_03420, two enzymes involved in the synthesis of methylerithriol phosphate. Finally, the three genes of the cluster (ERGA_CDS_03380, ERGA_CDS_03390, and ERGA_CDS_03400) have high ΔGC content and are located in a gene-sparse region of the genome. Altogether, this suggests that the ERGA_CDS_03390 cluster could be a functional cassette involved in *E. ruminantium* virulence and may have been acquired by horizontal gene transfer (HGT).

The two other conserved families of HATPase_c-containing effectors are exemplified by ERGA_CDS_02450, which encodes the heat shock protein 90 (HSP90), and ERGA_CDS_04380 *gyrB*, which encodes the beta subunit of DNA gyrase.

The HSP90 is an important cofactor in the response to oxidative stress and has been shown to have a crucial function in innate immune response (Mayor et al., 2007). This family of pT4Es may therefore be important for *Ehrlichia* to manipulate host immunity, possibly by avoiding or delaying pathogen recognition.

GyrB is a topoisomerase that is necessary for relaxation of DNA during replication. GyrB family of *Ehrlichia* pT4Es is expected to act in the nucleus and indeed it is noteworthy that they harbour NLS directly upstream from the HATPase_c domain. Controlling the state of DNA topology could be an efficient way for the bacterium to regulate specific host promoters. These proteins could thus mimic host chromatin-regulatory factors, as described for other effectors (Bierne and Cossart, 2012). The property of subtle mimicry of the activities of cellular proteins is also one of the most common features of type III secreted effectors (Galán, 2009). Moreover, a remarkable feature to note is that *Bartonella* toxin VbhT is secreted into target cells in a type IV-dependent manner and acts as a gyrase inhibitor. This VbhT effector represents the missing link in the evolution of *Bartonella* effectors from inter-bacterial conjugative toxins to inter-kingdom host-targeted effectors (Harms et al., 2017).

Similarly, the family of *Ehrlichia* HATPase_c-containing pT4Es could be a further example of the wonderful functional plasticity of effector proteins. It is worth mentioning that ERGA_CDS_04380 has been shown to be overexpressed in the virulent Gardel strain of *E. ruminantium* at early stages of infection (Emboulé, 2010).This is reminiscent of the early expression of *Ehrlichia* T4SS (Cheng et al., 2008) and compatible with T4SS-dependent secretion into the host cell. Thus, unravelling the role of HATPase_c-containing effectors in *Ehrlichia* pathogenesis will be a major challenge but could also highlight the importance of chromatin remodelling in *Ehrlichia* infections.

In this study, we showed that genomic plasticity *sensu largo, i.e*. from gene presence/absence polymorphisms to protein domain shuffling, with hallmarks such as local gene density, G+C content, intergenic recombination, is a major driver for *Ehrlichia* spp. to acquire and evolve new potential virulence functions. Our study revealed that *Ehrlichia* T4Es repertoires comprise core and variable gene suites, which likely have distinct roles in pathogenicity as well as different evolutionary histories. We also identified several DNA rearrangements in T4E genes, some of which could be correlated with the host specificity of *Ehrlichia* species. Despite the fact that more genomic sequences of *Ehrlichia* strains and functional validations are needed to provide evidence for robust associations between host range and T4E repertoires, our observations already suggest that the host range may be controlled by multiple or differential combinations of T4E determinants, or determinants other than T4E, or that differences in the T4E protein amino acids sequence (and even expression) may also be involved. We also identified new protein domain associations among *Ehrlichia* pT4Es, probably rearranged over the course of evolution, which could contribute to the numerous potential functions exerted by *Ehrlichia* effectors. A noticeable feature *of Ehrlichia* effectors is their modular architecture, comprising domains or motifs that could confer an array of subversive functions within eukaryotic cells. These domains/motifs therefore represent a fascinating repertoire of molecular determinants with important roles during infection.

Our study illustrates the power of comparative genomics to decipher the genetic basis of host specificity of this important family of pathogens. The properties in the cytoplasmic effector repertoires of *Ehrlichia* may predict the basis for pathogen host specificity. The study of these T4Es will provide insights not only into fundamental aspects of host-pathogen interactions, but also into the basic biology of eukaryotic cells.

Although the underlying basis for host specialization remains largely unresolved, adaption to a host clearly requires coevolutionary maintenance of a compatible effector repertoire. Further experimental determination of minimal *bona fide* T4Es repertoires required for *Ehrlichia* to cause to disease in a given host is needed. A major challenge in the future will be using systems-level knowledge in pathogen genomics, for different host-bacteria interactions, to predict the potential threat of emerging pathogens or to imagine science-based rapid response plans.

## Acknowledgements

This study was partly conducted in the framework of the project MALIN “Surveillance, diagnosis, control and impact of infectious diseases of humans, animals and plants in tropical islands” supported by the European Union in the framework of the European Regional Development Fund (ERDF) and the Regional Council of Guadeloupe. We warmly thank E. Albina and D. Goodfellow for reading and editing of this manuscript.

---

**Figure S1.**
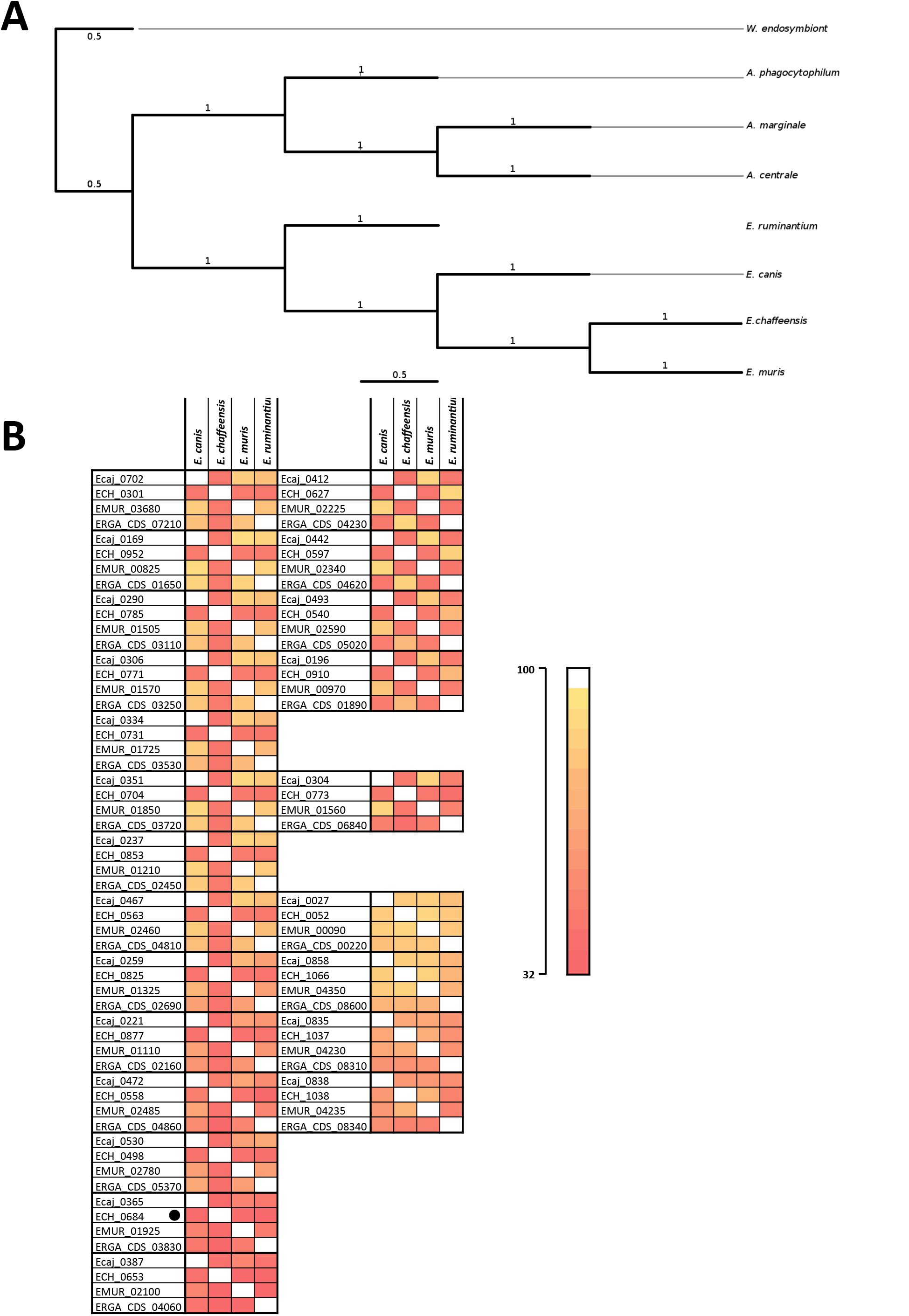
Phylogenetic tree of *Ehrlichia* and *Anaplasma* T4Es shows three different clades. **A.** A maximum-likelihood tree of 4 *Ehrlichia* species, 3 *Anaplasma* species and *W. endosymbiont* of *D. melanogaster* (out group) was reconstructed on the basis of concatenated nucleic acid alignments of pT4Es shared by all species (core effectome) with 100 bootstrap resamplings. **B.** The identity percentage was calculated for each effector ortholog group (EOG) of the *Ehrlichia* core effectome, and is represented by a heat map. The colour gradient represents the identity between effectors (pale colours mean high similarity).

**Figure S2.**
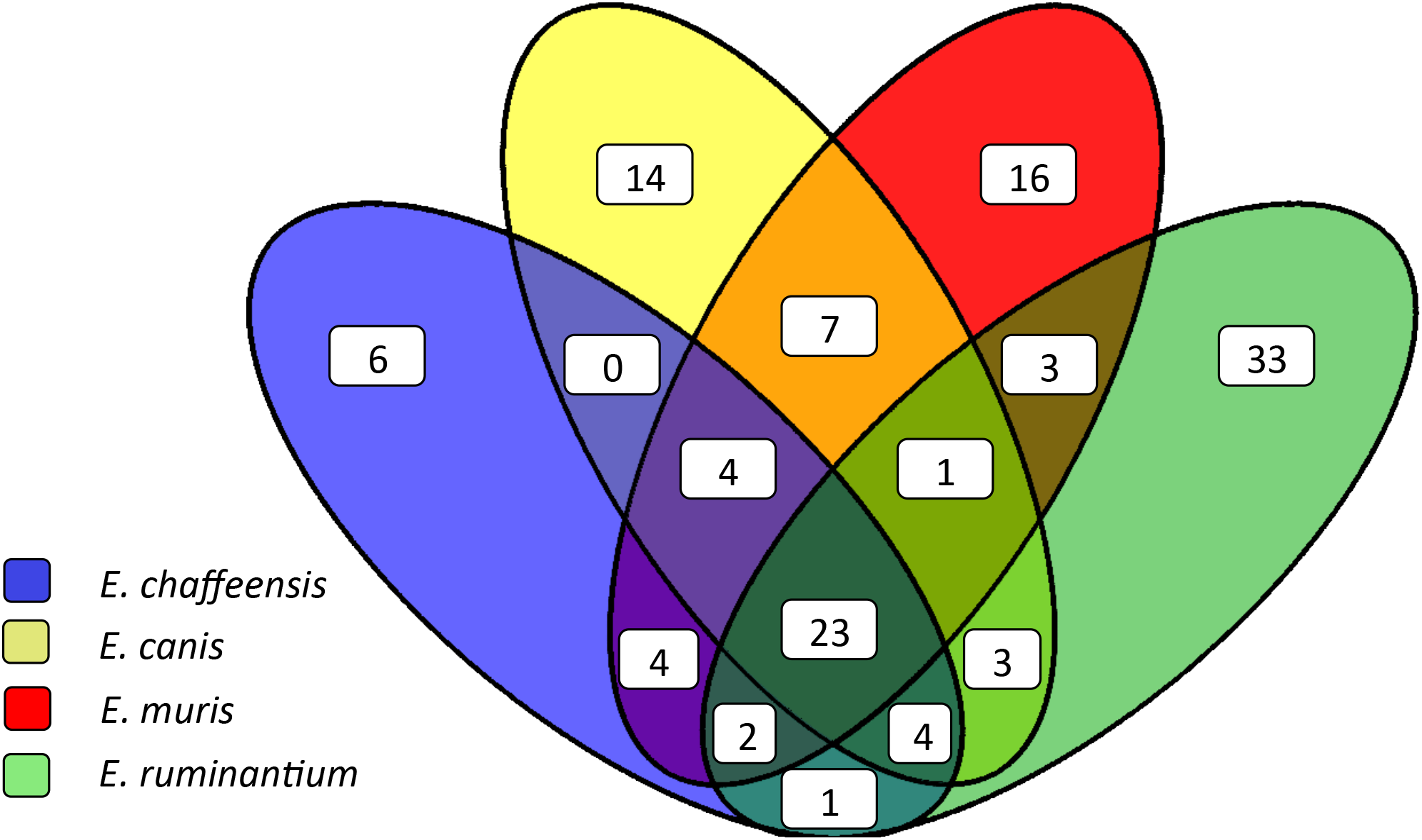
Venn diagram of *Ehrlichia* spp. pT4Es shows a large number of specific effectors. Predicted T4 effectomes of four *Ehrlichia* species compared with S4TE-CG and PanOCT to find homologous proteins in each species. Results are plotted on a Venn diagram and a number indicates the occurrence of predicted effectors is each intersection.

**Figure S3.**
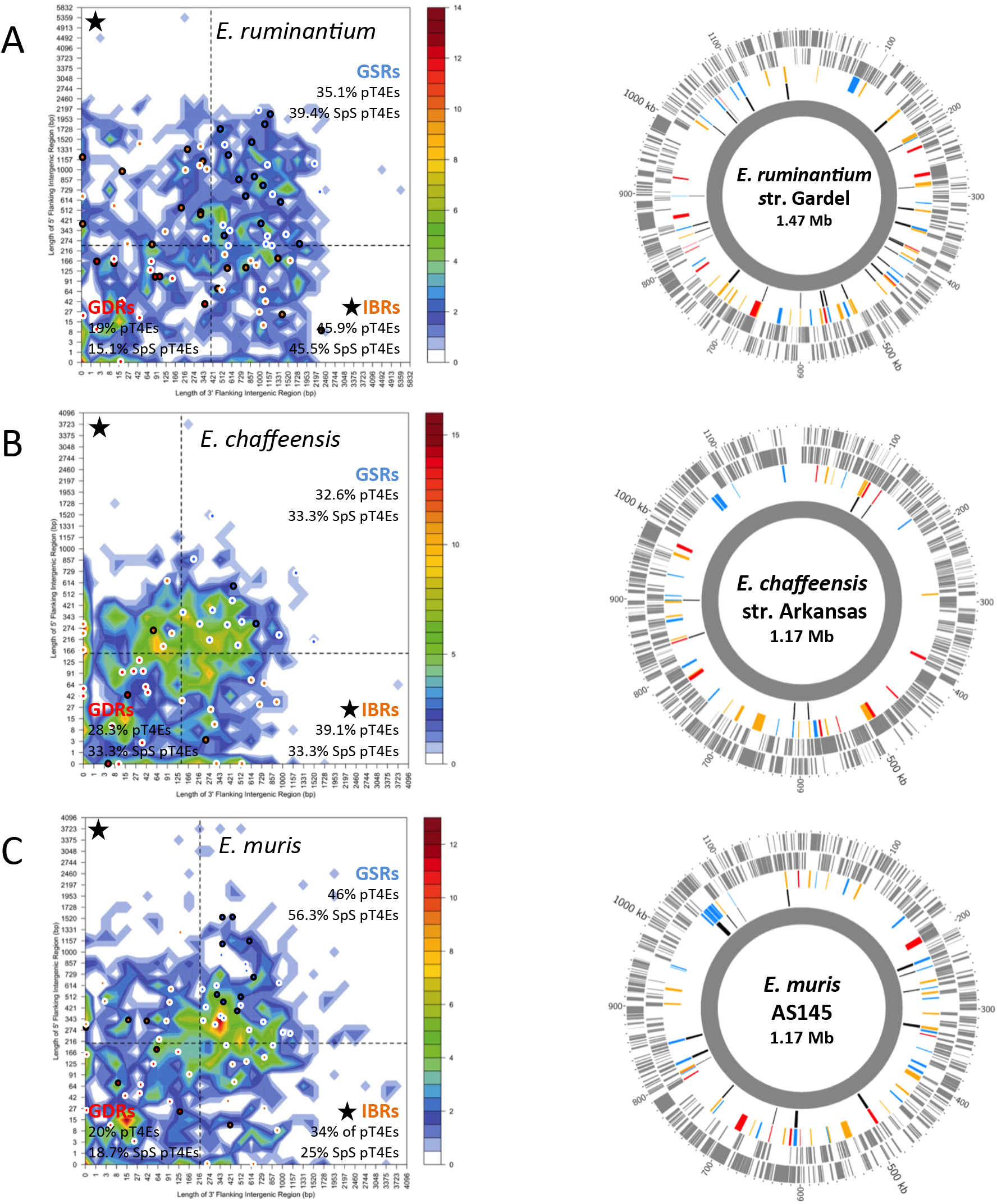
Distribution of predicted type IV effectomes according to local gene density based on the length of flanking intergenic regions (FIRs). Distribution of *E. ruminantium str*. Gardel, *E. chaffeensis* str. Arkansans and *E. muris* AS145 genes according to the length of their flanking intergenic regions (FIRs). All the genes of each species were sorted into two-dimensional bins according to the length of their 5⍰ (*y*-axis) and 3⍰ (*x*-axis) FIR lengths. The number of genes in the bins is represented by a colour-coded density graph. Genes whose FIRs were both longer than the median length of FIRs were considered as gene-sparse region (GSR) genes. Genes whose FIRs were both below the median value were considered as gene-dense region (GDR) genes. In between (IBR) genes are genes with a long 5⍰ FIR and short 3⍰ FIR, and inversely. For *E. ruminantium, E. chaffeensis* and *E. muris*, median values are 246 bp, 156 bp and 207 bp for 5⍰ FIRs respectively and 405 bp, 138 bp and 219 bp for 3⍰ FIRs respectively. The dashed line stands for the median length of FIR and delimits the genes in GSR, GDR and IBR. Candidate effectors predicted using the S4TE 2.0 algorithm were *s* plotted on this distribution according to their own 3⍰ and 5⍰ FIRs. A colour was assigned to each of the three following groups: red to GDRs, orange to IBRs, and blue to GSRs. Specific pT4Es are represented with a dot outlined in black. On the right, a Circos graph shows the distribution of *E. ruminantium* str. Gardel, *E. chaffeensis* str. Arkansans and *E. muris* AS145 pT4Es along the genome. The colour (red, orange or blue) of each gene corresponds to their location in GDR, IBR or GSR regions respectively.

**Figure S4.**
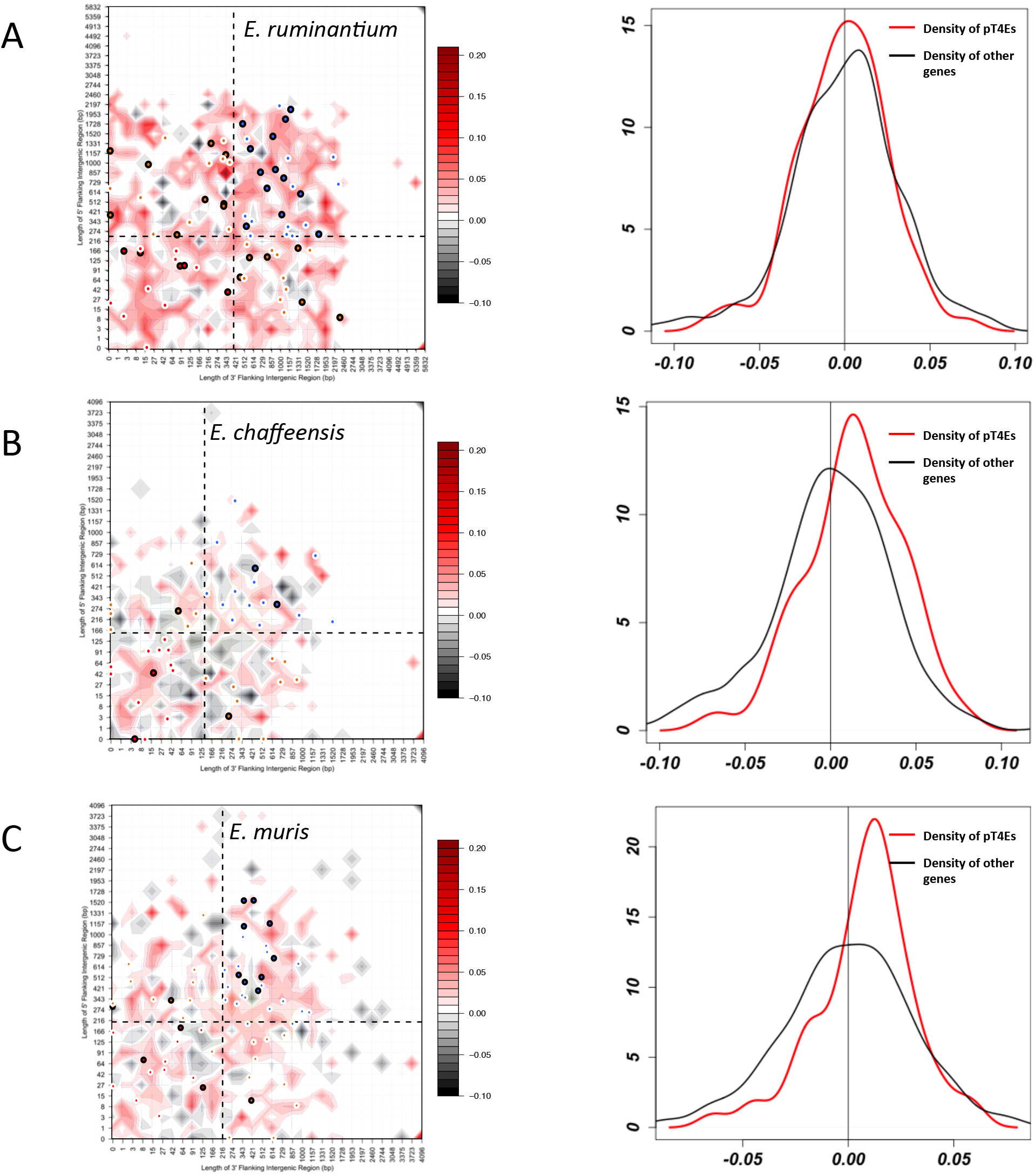
Distribution of the effectome according to the length of their flanking intergenic region (FIR) and ΔGC content. Distribution of *E. ruminantium* str. Gardel, *E. chaffeensis* str. Arkansans and *E. muris* AS145 genes according to the length of their flanking intergenic regions (FIRs). All the genes of each species were sorted into two-dimensional bins according to the length of their 5⍰ (*y*-axis) and 3⍰ (*x*-axis) FIRs. For each gene, the ΔGC content was calculated by subtracting the GC content of a gene by the average of GC content of all the genes. The mean of ΔGC of genes in the bins is represented by a colour-coded density graph. Genes whose FIRs were both longer than the median length of FIRs were considered as gene-sparse region (GSR) genes. Genes whose FIRs were both below the median value were considered as gene-dense region (GDR) genes. In between (IBR) genes are genes with a long 5⍰ FIR and short 3⍰ FIR, and inversely. For *E. ruminantium, E. chaffeensis* and *E. muris*, median values are 246 bp, 156 bp and 207 bp for 5⍰ FIRs, respectively, and 405 bp, 138 bp and 219 bp for 3⍰ FIRs, respectively. The dashed line showing the median length of FIR delimits the genes in GSR, GDR and IBR. A colour was assigned to each of the three following groups: red to GDRs, orange to IBRs, and blue to GSRs. Specific pT4Es are represented with a dot outlined in black. A density graph is plotted in the top right corner. The red line represents the density of pT4Es according to ΔGC content and the black line represents the density of the other genes.

